# Substrate-directed control of N-glycosylation in the endoplasmic reticulum calibrates signal reception at the cell-surface

**DOI:** 10.1101/2024.04.25.591210

**Authors:** Mengxiao Ma, Ramin Dubey, Annie Jen, Ganesh V. Pusapati, Evgenia Shishkova, Katherine A. Overmyer, Valérie Cormier-Daire, L. Aravind, Joshua J. Coon, Rajat Rohatgi

## Abstract

One-fifth of human proteins are N-glycosylated in the endoplasmic reticulum (ER) by two oligosaccharyltransferases, OST-A and OST-B. Contrary to the prevailing view of N-glycosylation as a housekeeping function, we identified an ER pathway that modulates the activity of OST-A. Genetic analyses linked OST-A to HSP90B1, an ER chaperone for membrane receptors, and CCCD134, an ER protein we identify as the first specificity factor for N-glycosylation. During its translocation into the ER, a N-terminal peptide in HSP90B1 functions as a pseudosubstrate inhibitor of OST-A and templates the assembly of specialized ER translocon complexes containing CCDC134. Unexpectedly, OST-A functions as a scaffold rather than an enzyme in this context, stabilizing HSP90B1 by preventing its hyperglycosylation and degradation. Disruption of this pathway impairs WNT signaling at the cell surface and causes the bone developmental disorder Osteogenesis Imperfecta. Thus, N-glycosylation can be regulated by ER factors to control cell-surface receptor signaling and tissue development.

**One-Sentence Summary:** N-glycosylation of asparagine residues on proteins can be regulated by specificity factors in the endoplasmic reticulum to control cell-surface signaling and tissue development.

## INTRODUCTION

Post-translational modifications are a central currency in signaling transactions because they allow protein activity to be regulated in response to changes in external signals or the internal state of the cell. For example, phosphorylation and ubiquitination on target proteins are tightly regulated by hundreds of substrate-specific kinases and E3 ligases, respectively. Asparagine-linked glycosylation or N-glycosylation in the endoplasmic reticulum (ER) is a pervasive post-translational modification in eukaryotes found on ∼20% of the proteome (*6*, *7*). Defects in N-glycosylation cause a class of human genetic diseases called congenital disorders of glycosylation (CDGs) affecting multiple organs, highlighting its importance in tissue development (*8*). In adult tissues, N-glycans participate in processes ranging from cell-cell recognition in the immune system, inflammatory responses and the metastatic spread of cancer cells (*1*, *9*). However, despite its myriad physiological and pathological roles, whether N-glycosylation can be a regulated post-translational modification (akin to phosphorylation) remains unknown.

The dominant view in cell biology is that N-glycosylation is a constitutive process initiated by two multi-subunit oligosaccharyltransferase complexes, OST-A and OST-B, that contain the distinct catalytic subunits STT3A and STT3B, respectively (*2*). OST-A, which contains unique subunits that anchor it to the SEC61 protein channel (*10*), functions co-translationally to transfer a pre-assembled glycan *en bloc* from a lipid donor to asparagine residues found in the context of N-X-S/T/C sequons (where X can be any residue other than proline). Structural studies show that the catalytic site in STT3A is positioned at the luminal surface of the ER membrane to scan the nascent chain for sequons as it emerges from the SEC61 translocon (*11–14*). OST-B functions co- or post-translationally on sequons missed or poorly recognized by OST-A (*2*). Sequons are enriched on luminal or extracellular domains of proteins and are predominantly located on surface-exposed loops and turns (*7*).

The branched Glc_3_Man_9_GlcNAc_2_ glycan (comprising 2 N-acetylglucosamine, 9 mannose, and 3 glucose molecules) installed by OST-A and OST-B is edited by various glycosyltransferases and glycosidases as proteins travel through the secretory pathway (*15*, *16*). In the ER, terminal glucose and mannose residues are cyclically removed and attached as part of a timing mechanism that triages proteins into protein folding, degradation and ER export pathways. In the Golgi, further editing of the core glycan generates a number of N-glycoforms, classified as high-mannose, hybrid and complex N-glycans, the latter of which serve multiple functions at the cell surface.

An important unanswered question is whether N-glycosylation can be regulated to regulate cellular processes. Global mass spectrometry data shows significant N-glycoside heterogeneity (termed ‘macroheterogeneity’) across different cells and tissues, suggesting that regulated N-glycosylation may represent a pervasive (and largely unexplored) layer of cellular regulation (*17*). Previous studies have found that the transcriptional regulation of genes encoding Golgi-localized glycosyltransferases, accessibility of sequons, availability of glycan substrates and trafficking itinerary of proteins can all influence the final complement of N-glycoforms presented on the cell surface (*16*, *18*). However, whether the activity of the OST-A and OST-B complexes, which sit at the apex of the N-glycosylation pathway, can be controlled by other factors to switch the glycan occupancy of specific sequons on or off remains unknown. In the course of a screen for new components of the WNT signaling pathway, we serendipitously discovered the first example of a signaling module that controls the activity of OST-A towards specific sequons on a specific protein. This pathway regulates the sensitivity to WNT ligands at the cell surface, providing a mechanism for communication between the ER and the plasma membrane.

## RESULTS

### Genetic screens identify an ER-localized regulator of WNT signaling

The WNT pathway is a highly conserved metazoan cell-cell communication system that guides both tissue development during embryogenesis and tissue homeostasis and repair in adults (*19*). Using a fluorescence-based transcriptional reporter of WNT signaling strength (*20*), we conducted a genome-wide, loss-of-function CRISPR/Cas9 screen in a human cell line (RKO) to identify genes required to activate WNT/β-catenin signaling in response to WNT3A (**Fig.1A** and **Data S1**). The screen identified many known WNT signaling components (**Fig.1B**): β-catenin (gene *CTNNB1)*, LRP6 (a co-receptor that binds WNT ligands in cooperation with Frizzled (FZD) proteins), and two proteins, MESD (gene *MESDC2*) and HSP90B1, that function as chaperones to facilitate LRP5/6 folding in the ER (*21*, *22*). Our screens also identified several genes not clearly linked to WNT signaling: *CCDC134*, *STT3A* and *OSTC*. STT3A and OSTC are two of the eight components of the OST-A complex. CCDC134 is a poorly studied protein named for the presence of a coiled-coil domain; however, structure prediction and sequence complexity analysis instead show that it contains a core globular domain composed of a unique 5-helix bundle (**fig.S1A**). CCDC134 is required for mouse embryonic development and some studies propose it is a secreted cytokine that attenuates the MAPK pathway (*23*, *24*). However, CRISPR/Cas9-mediated disruption of *CCDC134* in multiple cell lines resulted in reduced responsiveness to WNT ligands, measured by WNT reporter activity or β-catenin protein accumulation (**Fig.1C,1D,1E**). A single study (*25*) suggested that CCDC134 is “possibly” linked to WNT signaling based on altered expression of some genes peripherally related to WNT signaling in the cerebellum; however, no evidence has been presented to show an effect on WNT signal transduction or WNT target genes.

**Figure 1.**
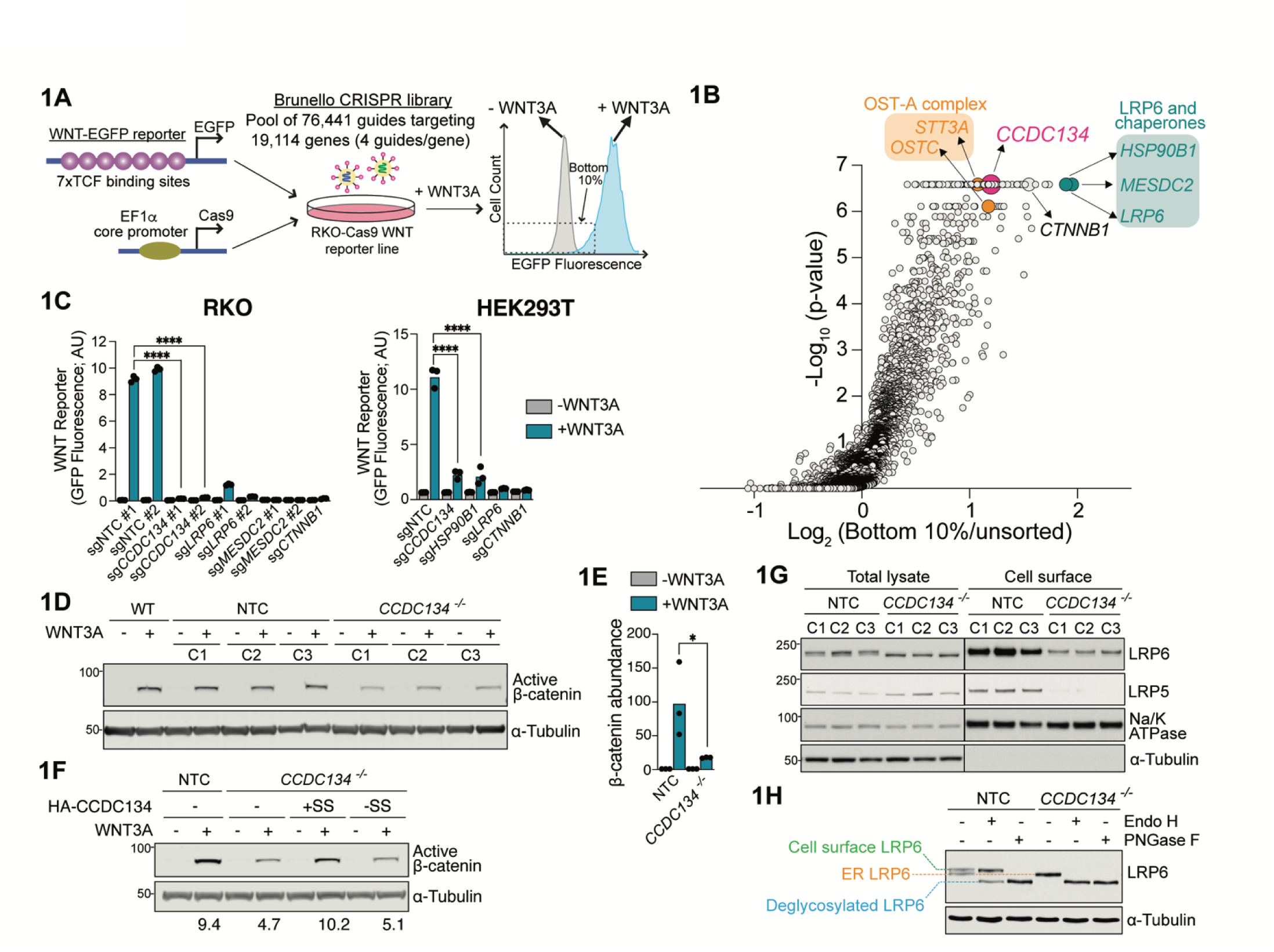
CCDC134 is an ER protein that promotes cell-surface abundance of the WNT co- receptors LRP5 and LRP6. **(A)** Screening strategy used to identify positive regulators of WNT signaling (see text for details). **(B)** Volcano plot of CRISPR/Cas9 screen results. Each circle is one gene, the x-axis shows its enrichment or depletion calculated as the mean of all four sgRNAs targeting the gene in the bottom 10% sorted population relative to the unsorted population, and the y-axis shows statistical significance as measured by *p*-value. Genes that are the focus of this study are highlighted in color. *CTNNB1*, which encodes β-catenin, is highlighted as a positive control. Full screen results are provided in **Data S1**. **(C)** WNT-EGFP reporter fluorescence (+/- WNT3A conditioned media treatment) in RKO (left) or HEK293T cells (right) expressing a non-targeting control sgRNA (NTC) or sgRNAs (different from those used for the CRISPR/Cas9 screen in **1B**) targeting selected screen hits from (**B**). Each data point represents median WNT-EGFP reporter fluorescence from one population and bars show the mean of three populations. Statistical significance was determined by one-way ANOVA Sidak’s multiple comparisons test; **** p<0.0001 (*n*=3, ∼3300 cells each). **(D, E)** Active (non-phosphorylated) β-catenin abundance (a metric of WNT signaling strength) was measured (+/- WNT3A) using immunoblots in wild-type cells or clonal cell lines expressing a control (NTC) sgRNA or an sgRNA targeting *CCDC134*. The graph in (**E**) shows active β-catenin abundance normalized to α-tubulin abundance in three independently derived clonal cell lines (C1-C3, shown in **D**), with the bar representing the mean. Statistical significance was determined by one-way ANOVA Dunnett’s multiple comparisons test; * p<0.05. (F) Active β-catenin abundance (+/- WNT3A) in clonally derived control (NTC) and *CCDC134^-/-^* cell lines stably expressing near endogenous levels of 3xHA-CCDC134 (see **fig.S1B**) carrying an N-terminal ER signal sequence (+SS) or, as a control, lacking a signal sequence (-SS) to prevent ER targeting. Numbers below the lanes show the WNT3A-induced fold-increase in active β-catenin abundance normalized to α-Tubulin abundance. (G) LRP6 and LRP5 abundances in total lysate (6% input) or the plasma membrane (cell surface biotinylation and streptavidin immunoprecipitation, 33% elution) from three independently derived (C1- C3) control (NTC) or *CCDC134^-/-^* clonal cell lines. The cell surface protein Na/K ATPase and cytoplasmic protein α-tubulin serve as controls, both for loading and for the specificity of cell surface biotinylation. The ER population and the cell surface population of LRP6 are visible as bands of slightly different mobilities on immunoblots (see **1H**). (H) Glycosidase sensitivity in conjunction with mobility on SDS-PAGE gels was used to measure the ER or cell-surface pools of LRP6 in control or *CCDC134^-/-^* cells. Endoglycosidase H (Endo H) can remove glycans added in the ER but not the complex glycan modifications added in the Golgi; Peptide-*N*- Glycosidase F (PNGase F) can remove glycans on both ER and cell-surface proteins. **See also fig.S1.**

In apparent contradiction to its annotation (*23*) as a protein secreted into the extracellular space, we noted that CCDC134 contained a ‘QSEL’ sequence at the extreme C-terminus, separated from the its helical globular domain by an unstructured linker (**fig.S1A**). This sequence motif is a variant of the canonical ‘KDEL’ sequence and is known to mark proteins that are retained in the ER, rather than secreted from the cell (*26*). Indeed, CCDC134 from other animal species contains a more canonical ER retention sequence (e.g., KEEL, KNEL, RDEL). For immunoblot and immunofluorescence analysis, we constructed a tagged form of CCDC134 with a 3xHA tag at the N-terminus after the ER signal sequence (**fig.S1A**) to avoid disrupting ER localization and retention. Expression of 3xHA-CCDC134 was titrated using doxycycline to achieve near endogenous protein abundance (**fig.S1B**). Our experiments confirmed that CCDC134 is an ER resident protein: mutation of its C-terminal QSEL to QSSA abrogated its ER retention and enabled its secretion into the media (**fig.S1C, S1D**). Importantly, the CCDC134-QSSA mutant failed to restore WNT signaling to wild-type levels when expressed in *CCDC134*^-/-^ cells (**fig.S1E**). CCDC134 is also annotated as a nuclear protein in some studies (*27*); however, it has a well-defined signal sequence for ER insertion and is not enriched in the nucleus (**fig.S1C**). CCDC134 expressed without an ER signal sequence failed to restore WNT signaling when expressed in *CCDC134*^-/-^ cells (**Fig.1F**). We conclude that CCDC134 is incorrectly annotated in the literature and in widely used databases like UniProt: it is not a secreted or nuclear protein, but rather functions in the ER.

During experiments designed to understand how CCDC134 regulates WNT signaling, we discovered that cell-surface LRP5 and LRP6 levels were markedly reduced in *CCDC134*^-/-^ cells (**Fig.1G**). Electrophoretic mobility and glycosylation analyses (*28*) (**Fig.1H**) revealed that LRP5/6 is trapped in the ER in *CCDC134*^-/-^ cells: LRP6 shows a subtle increase in its electrophoretic mobility (seen by a downward shift on gels, **Fig.1G**), typically observed when membrane proteins are trapped in the ER and so fail to acquire the complex glycans added in the Golgi *en route* to the cell surface. Consistent with ER retention, LRP6 in *CCDC134*^-/-^ cells was completely sensitive to Endo H (**Fig.1H**), a glycosidase that can remove the N-glycans found on proteins in the ER but cannot remove the complex glycans found on cell surface proteins (*29*). In contrast, both the ER (Endo H sensitive) and cell-surface (Endo H resistant, PNGase F sensitive) populations of LRP6 were observed in wild-type cells as two bands of distinct mobility on SDS-PAGE gels (**Fig.1H**). In summary, CCDC134 regulates WNT signaling in the ER by controlling the trafficking of LRP5/6, obligate co-receptors for WNT ligands, to the cell surface.

### CCDC134 regulates the stability of an ER chaperone for the WNT co-receptors LRP5/6

The abundance of LRP5/6 on the cell surface depends on MESD and HSP90B1. Only correctly folded LRP5/6 is allowed to move from the ER to the Golgi and eventually to the cell surface by the ER quality control system. The disruption of either HSP90B1 or MESD leads to the depletion of LRP5/6 from the cell surface (*21*, *22*), reminiscent of the *CCDC134*^-/-^ phenotype. Immunoblotting (**Fig.2A**) and mass spectrometry (**Fig.2B**) both revealed that HSP90B1 levels were reduced in *CCDC134*^-/-^ cells, but MESD levels were unchanged (**Fig.2A**). Additionally, a sub-population of HSP90B1 migrated as a higher molecular weight species on SDS-PAGE gels in *CCDC134*^-/-^ cells compared to wild-type cells (**Fig.2A** and **fig.S2A**). After digestion with Endo H or PNGase F, the HSP90B1 band from both wild-type and *CCDC134*^-/-^ cells migrated at the same position, suggesting that HSP90B1 acquired additional N-glycans in *CCDC134*^-/-^ cells (**Fig.2C**). Rescue experiments demonstrated that re-expression of CCDC134 both suppressed HSP90B1 glycosylation and restored its abundance (**Fig.2D**).

**Figure 2.**
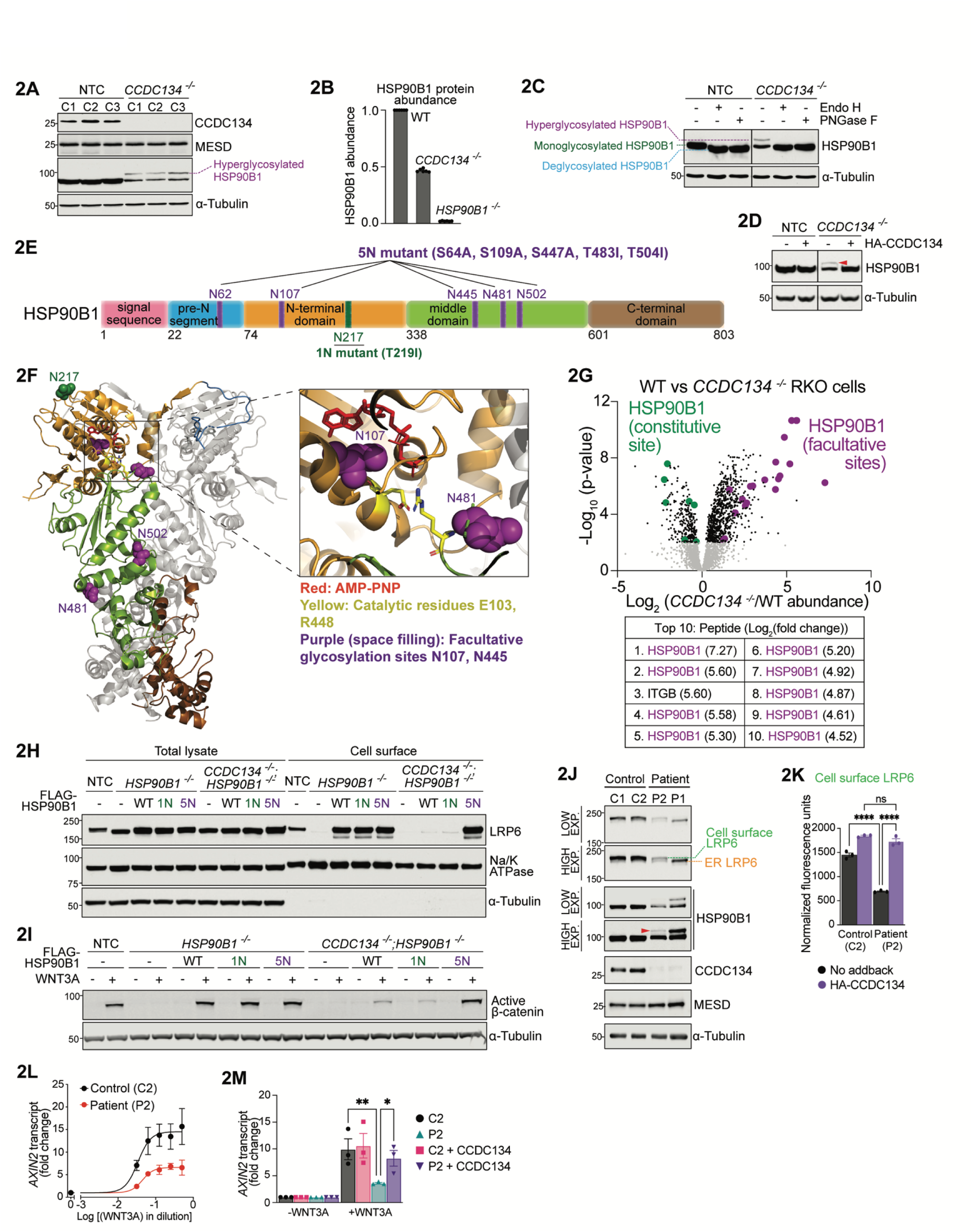
Hyperglycosylation of HSP90B1 regulates WNT signaling strength. **(A)** Abundance and glycosylation status of HSP90B1 on SDS-PAGE gels in lysates from three (C1-C3) independent control (NTC) and *CCDC134^-/-^*clonal cell lines. **(B)** HSP90B1 protein abundance in wild-type (WT), *CCDC134^-/-^*, *HSP90B1^-/-^* RKO cell lines measured by mass spectrometry, normalized to WT cells. Bars represent the mean abundance +/- SEM from six independent mass spectrometry runs, represented as individual data points. **(C)** Glycosidase sensitivity (Endo H or PNGase F see Fig.1H) of HSP90B1 in lysates of control (NTC) or *CCDC134^-/-^* cells. HSP90B1 is glycosylated at either a single constitutive site (monoglycosylated band) or at multiple facultative sites (hyperglycosylated bands) (see (*30*) and text for details). **(D)** Abundance and glycosylation status of HSP90B1 (hyperglycosylated form is denoted by a red arrowhead) in lysates from control (NTC) or *CCDC134^-/-^* cell lines with or without CCDC134 expression from a doxycycline-inducible, stably integrated transgene (see **fig.S1B**). **(E)** Domain architecture of HSP90B1 as defined in(*58*). The constitutive and facultative N-glycosylation sites are labeled in green and purple, respectively. The point mutations used to disrupt the one constitutive sequon (1N mutant) and the five facultative sequons (5N mutant) are listed. The N-terminal domain (NTD) binds ATP, the Middle domain (MD) participates in ATP hydrolysis and client recognition, and the C-terminal domain (CTD) in dimerization. **(F)** Structure (PDB 5ULS) highlighting one protomer of a HSP90B1 dimer, with the pre-N, NTD, MD and CTD domains colored as in (**E**) (*58*). The ATP bound to the NTD and the asparagine side chains of the constitutive sequon (green) and the facultative sequons (purple) are shown in space filling representation. The inset shows proximity of two facultative sequons to the ATP binding site. **(G)** Abundance of glycosylated peptides in *CCDC134^-/-^* cells measured using global, unbiased N- glycoprotemics. Each data point (top graph) represents the fold change in abundance of a distinct glycopeptide (defined by sequence and glycan structure) in *CCDC134^-/-^* compared to wild-type cells. Glycopeptides that include the constitutive and facultative sequons in HSP90B1 are colored green and purple, respectively. Amongst the ten glycopeptides (in the entire proteome) that show the greatest fold- increase in *CCDC134^-/-^* cells, nine are from regions of HSP90B1 that include the facultative sites (bottom Table). Full dataset is provided in **Data S2**. **(H, I)** Abundances of cell-surface LRP6 (**H**) or active β-catenin abundance (**I**) in clonally derived control (NTC), *HSP90B1^-/-^*, and *CCDC134^-/-^*;*HSP90B1^-/-^* cell lines stably expressing wild-type (WT) FLAG- HSP90B1 or variants carrying mutations in the one constitutive (1N) or all five facultative (5N) sites (see Fig.2E). Abundances of stably expressed HSP90B1 variants were comparable (**fig.S2E**). **(J-M)** LRP6 and HSP90B1 abundances (**J**), WNT3A dose-response curves (**L**), WNT signaling strength (**M**), and cell-surface LRP6 levels measured by flow-cytometry (**K**) in primary fibroblasts isolated from two OI patients carrying homozygous loss-of-function mutations in CCDC134 (c.2T>C) (P1, P2) and two age-matched healthy control individuals (C1 and C2, respectively)(*5*). (**L,M**) The abundance of *AXIN2* mRNA, encoded by an immediate-early WNT target gene, was measured by quantitative reverse transcription PCR (qRT-PCR) as a metric of WNT signaling strength. In **K**, **L** and **M** error bars show the mean +/- SEM from three independent experiments. Statistical significance was determined by two-way ANOVA Dunnett’s multiple comparisons test; * p<0.05, ** p<0.01 (**M**) or two-way ANOVA Tukey’s multiple comparisons test; **** p<0.0001 (**K**). **See also fig.S2.**

The excessive N-glycosylation (hereafter called “hyperglycosylation”) of HSP90B1 has been observed previously, but its function and regulation have remained mysterious for three decades (*4*, *30*–*33*). In these prior studies, hyperglycosylation was observed when HSP90B1 was overexpressed or when cells were subjected to ER stress. Under each of these conditions, co-expression of CCDC134 was able to suppress HSP90B1 hyperglycosylation (**fig.S2B, S2C**). HSP90B1 has six glycosylation sites conserved in evolution (*32*) (**Fig.2E**). One of these sequons (N217) is constitutively modified with an N-glycan (hereafter called the “constitutive sequon”) while five (N62, N107, N445, N481 and N502) are only used under conditions where HSP90B1 is hyperglycosylated (hereafter called “facultative sequons”) (*4*, *30*, *32*). The facultative sequons are located in regions of HSP90B1 that would be predicted to impair protein function or folding (*4*) (**Fig.2F**). This arrangement is unusual: in most secretory pathway proteins, sequons have been depleted during evolution from regions (e.g. the buried interior or at active sites) where N-glycosylation would be expected to disrupt folding or function (*7*, *34*). Hyperglycosylated HSP90B1 likely cannot fold into a functional protein, causing it to be flagged and degraded by the ERAD machinery (*4*). Accordingly, hyperglycosylated HSP90B1 is stabilized by two structurally unrelated inhibitors of the ERAD pathway (**fig.S2D**).

To measure the effect of CCDC134 loss on global N-glycosylation, we used a specialized mass spectrometry platform(*17*) for the unbiased, proteome-wide identification of N-glycosylated peptides from wild-type and *CCDC134*^-/-^ cells. N-glycoprotemics revealed a strikingly specific effect of CCDC134 on the glycosylation of the facultative sequons in HSP90B1. Out of almost 4000 identified N-glycopeptides, those derived from HSP90B1 showed the greatest increase in abundance in *CCDC134*^-/-^ cells compared to wild-type cells (**Fig.2G**), despite the fact that the overall abundance of HSP90B1 decreased (**Fig.2A,2B**).

To test whether hyperglycosylation of HSP90B1 was the root cause of reduced cell-surface LRP5/6 and WNT sensitivity in *CCDC134*^-/-^ cells, we disrupted all five facultative sequons in HSP90B1 (hereafter called the HSP90B1^5N^ mutant) (**Fig.2E**) (*32*). As a control, we also separately disrupted the single constitutive sequon (N217, mutated in HSP90B1^1N^). Both HSP90B1^5N^ and HSP90B1^1N^ were functional proteins because they were expressed at levels comparable to wild-type HSP90B1 (**fig.S2E**) and could rescue cell surface LRP6 and WNT signaling in *HSP90B1*^-/-^ cells (**Fig.2H**). However, only HSP90B1^5N^ (but not wild-type HSP90B1 or HSP90B1^1N^) could rescue *HSP90B1*^-/-^;*CCDC134*^-/-^ double null cells (**Fig.2H,2I**), formally showing that HSP90B1 is epistatic to CCDC134 in the WNT pathway. These data provide evidence that HSP90B1 hyperglycosylation is the cause of impaired WNT signaling in *CCDC134*^-/-^ cells (rather than merely a correlated observation). A parsimonious model supported by these data is that the loss of CCDC134 leads to the hyperglycosylation and ERAD-mediated degradation of HSP90B1. When ER levels of HSP90B1 drop, LRP5/6 folding and subsequent cell surface trafficking is impaired, reducing sensitivity to WNT ligands.

### HSP90B1 hyperglycosylation and LRP6 trafficking in human patients carrying *CCDC134* mutations

This role of CCDC134 in regulating HSP90B1 is not restricted to the RKO cells in which we performed the original CRISPR/Cas9 screen. First, loss of CCDC134 led to HSP90B1 hyperglycosylation in a panel of other mouse and human cell lines (**fig.S2F**). Second, human genetic studies recently identified five patients (from three independent families) afflicted with the bone developmental disorder Osteogenesis Imperfecta (OI) who carry loss-of-function mutations in the *CCDC134* gene (*5*, *35*, *36*).

These patients suffer from a severe, deforming subtype of OI (Type III) that is also intriguingly seen in patients carrying mutations in other WNT pathway genes such as *MESDC2* and *WNT1* (*37–42*). A large body of evidence from both mouse and human studies has shown that WNT signaling, particularly its LRP5/6 node (*43–45*), plays an instructive role in the specification, differentiation and maintenance of osteoblasts. WNT/β-catenin signaling regulates bone development by driving progenitors along the osteoblast lineage to produce mature osteoblasts with a high capacity to synthesize components of the bone matrix (*46–49*). *CCDC134*^-/-^ osteoblasts isolated from patients show a defect in this same WNT-dependent step-- the ability to differentiate into highly secretory mature osteoblasts (*5*).

We tested the consequences of CCDC134 loss on primary fibroblasts isolated from two different *CCDC134*^-/-^ patients (**Fig.2J-2M** and **fig.S2G-S2I**). Consistent with our findings in cultured cells, extracts from patient fibroblasts showed reduced HSP90B1 abundance accompanied by evidence for hyperglycosylation (**Fig.2J**). Total and cell-surface LRP6 was reduced (**Fig.2J,2K)** and responses to WNT3A were blunted (**Fig.2L** and **fig.S2G**). Stable re-expression of CCDC134 in patient-derived fibroblasts using lentiviral delivery suppressed HSP90B1 hyperglycosylation (**fig.S2I**) and restored both cell-surface LRP6 (**Fig.2K**) and WNT signaling (**Fig.2M** and **fig.S2H**). Thus, our mechanistic insights into how CCDC134 influences WNT signaling were confirmed in primary human fibroblasts derived from *CCDC134*^-/-^ OI patients, establishing organism-level functional and disease relevance.

### Genetic interaction analysis links CCDC134 to the oligosaccharyltransferase OST-A

Previous work has shown that the knockout of STT3A, the catalytic subunit of the OST-A complex, but not STT3B results in hyperglycosylation of HSP90B1, mimicking the consequences of CCDC134 loss (*30*). Genes encoding two components exclusive to the OST-A complex, STT3A and OSTC, were also amongst the most significant positive regulators of WNT signaling in our CRISPR/Cas9 screen (**Fig.1B**). Finally, CCDC134, HSP90B1, STT3A and OSTC show a strong signature of genetic co-essentiality in the Cancer Dependency Map (DepMap) (**Fig.3A**) (*50*). Co-essentiality implies that they may function in the same biochemical pathway because they have concordant effects on fitness in loss-of-function CRISPR/Cas9 screens performed across hundreds of cancer cell lines. These three orthogonal observations suggested that CCDC134 may regulate N-glycosylation of HSP90B1 by regulating the OST-A complex.

**Figure 3.**
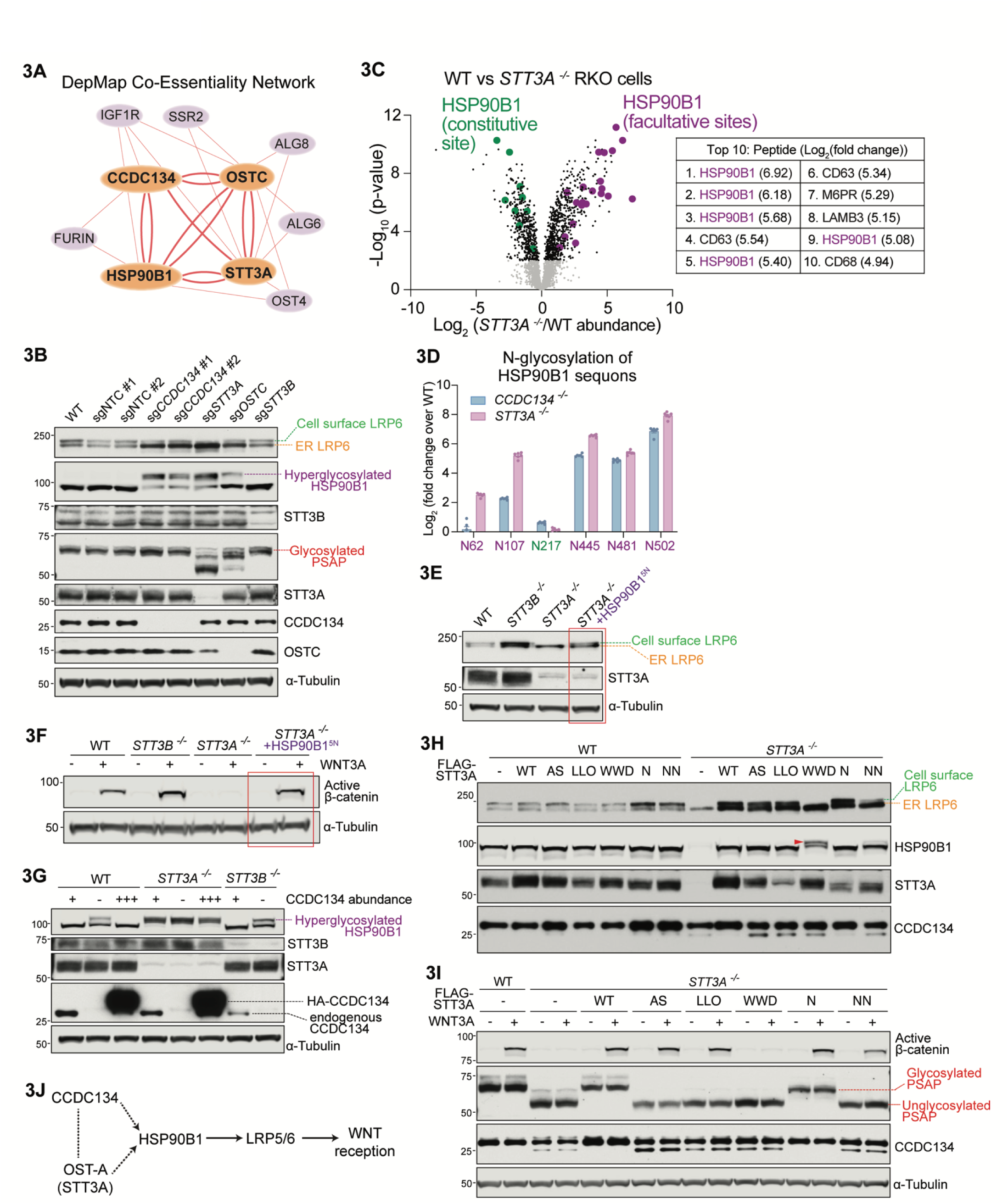
Genetic interactions between CCDC134, HSP90B1 and the OST-A complex. **(A)** DepMap co-essentiality relationships between CCDC134, STT3A, HSP90B1, and OSTC visualized using *Fireworks* (*70*). The bi-directional edges between these four genes (bold orange lines) indicate concordant effects on cell growth across >400 cell lines. **(B)** Abundances and glycosylation status of the indicated proteins (labeled to the right of each immunoblot segment) in lysates from cells expressing a control (NTC) sgRNA or sgRNA’s against various genes. **(C)** Abundance of glycosylated peptides in *STT3A^-/-^* cells compared to WT cells (represented as Fig.2G). Glycopeptides that include the constitutive and facultative sequons in HSP90B1 are colored green and purple, respectively. Amongst the ten glycopeptides in the entire proteome that show the greatest fold- increase in *STT3A^-/-^*cells, five are from regions of HSP90B1 that include the facultative sites (table on right). Full dataset is provided in **Data S2**. **(D)** Enrichment of glycopeptides (normalized to total HSP90B1 protein abundance from mass spectrometry) that include each of the facultative and constitutive sequons of HSP90B1 in *CCDC134^-/-^* or *STT3A^-/-^* cells compared to WT cells. Bars show the mean +/- SEM from six independent mass spectrometry runs, represented as individual data points. The abundances of all peptides that include each HSP90B1 sequon were integrated using individual peptide data summarized in Fig.2G and **3C**). (**E, F**) Abundances of LRP6 (**E**) and active β-catenin (**F**) in WT or clonal *STT3B^-/-^* and *STT3A^-/-^* cells, with or without stable expression of a variant of HSP90B1 (HSP90B1^5N^, Fig.2E) carrying disrupting mutations in all five facultative sequons. (**G**) Abundance and glycosylation status of HSP90B1 in *STT3B^-/-^*and *STT3A^-/-^* cells expressing different levels of CCDC134: No CCDC134 (-), endogenous CCDC134(**+**), stably overexpressed 3xHA-CCDC134 (**+++**) on top of endogenous CCDC134. (**H,I**) Abundances of LRP6 and HSP90B1 (**H**) or active β-catenin (**I**) in WT or *STT3A^-/-^* cells stably expressing FLAG-STT3A variants carrying mutations in various sites involved in catalytic transfer of the glycan from the lipid-linked oligosaccharide to the asparagine in sequons. Variants (shown on a structure in **fig.S3A**) carry mutations in residues involved in active site (AS) chemistry, lipid-linked oligosaccharide binding (LLO), sequon binding (WWD) or N-glycosylation of STT3A itself (N and NN). Glycosylation of PSAP (**I**) was used to assess OST-A activity in cells. CCDC134 serves as a loading control in (**H**). (**J**) A provisional pathway diagram constructed based on genetic interactions (dotted lines) uncovered in Fig.2 and 3 and physical interactions (solid lines) described in the literature. See also **fig.S3.**

The CRISPR/Cas9-mediated disruption of STT3A or OSTC, but not STT3B, led to HSP90B1 hyperglycosylation and a consequent reduction in cell-surface LRP6 (**Fig.3B**). As previously reported in HEK293 cells (*30*), global N-glycoproteomics demonstrated that N-glycopeptides derived from the facultative sequons in HSP90B1 showed the highest increase in abundance in *STT3A*^-/-^ cells compared to wild-type cells (**Fig.3C**). The abundance of glycopeptides (corrected for total HSP90B1 protein abundance) encompassing four out of the five facultative HSP90B1 sequons increased concordantly in *STT3A*^-/-^ and *CCDC134*^-/-^ compared to wild-type cells (**Fig.3D**). As expected, the abundance of glycopeptides containing the constitutive sequon (N217) did not change.

Analogous to the loss of either CCDC134 or HSP90B1 (**Fig.1D,1E,1G-I**), disruption of STT3A also markedly reduced cell-surface LRP6 and the strength of WNT signaling (**Fig.3E,3F**). Global disruption of N-glycosylation in *STT3A*^-/-^ cells could impact WNT signaling through multiple indirect mechanisms, such as hypoglycosylation of membrane receptors or activation of the unfolded protein response (*30*, *51*). To test whether reduced WNT signaling in *STT3A*^-/-^ cells is caused specifically by disruption of the CCDC134-HSP90B1 pathway, we took advantage of the HSP90B1^5N^ mutant, which can bypass the requirement for CCDC134 in WNT signaling (**Fig.2I**). Strikingly, HSP90B1^5N^ expression was also sufficient to rescue WNT signaling in *STT3A*^-/-^ cells (**Fig.3E,3F**), despite the fact that the N- glycosylation of hundreds of membrane and secreted proteins is altered when OST-A function is lost (*30*). CCDC134 overexpression can suppress HSP90B1 hyperglycosylation in many contexts (**fig.S2B,S2C**); however, even a massive increase in CCDC134 abundance was unable to suppress HSP90B1 hyperglycosylation in *STT3A*^-/-^ cells (**Fig.3G**).

Several key residues in STT3A have been identified as being important for its function (summarized in (*52*)). These are involved in binding to the lipid-linked oligosaccharide (LLO) (*53*, *54*), binding to the N-X-S/T sequon (*55*), catalyzing the chemical step in LLO transfer to the carboxamide side chain of asparagine (active site residues) (*56*) and in N-glycosylation of STT3A itself (*57*) **(fig.S3A**). We stably expressed STT3A proteins carrying established mutations in *STT3A*^-/-^ cells and assessed N- glycosylation of both HSP90B1 and prosaposin (PSAP). The latter is a substrate exclusively glycosylated by OST-A (*2*, *51*) and hence useful as a metric for its enzymatic activity in cells. Unexpectedly, inactive STT3A variants carrying mutations in the active site or LLO binding site were both fully able suppress HSP90B1 hyperglycosylation (unlike *STT3A*^-/-^ cells), restore cell-surface levels of LRP6 (**Fig.3H**), and rescue WNT signaling (**Fig. 3I**). However, the STT3A variant carrying mutations in the sequon binding site (WWD) was clearly defective in suppressing HSP90B1 hyperglycosylation. We conclude that the oligosaccharyltransferase activity of OST-A is dispensable for its ability to regulate HSP90B1 hyperglycosylation. Instead, OST-A may function as a translocon-proximal scaffold to suppress HSP90B1 hyperglycosylation in a manner that requires its sequon binding site. Indeed, disruption of OSTC (the subunit that anchors OST-A to the translocon) in cells expressing only the catalytically inactive, active site mutant of STT3A also triggered HSP90B1 hyperglycosylation (**fig.S3B**).

The ability of the HSP90B1^5N^ mutant to bypass the loss of CCDC134 (**Fig.2H,2I**) or STT3A (**Fig.3E,3F**) places *HSP90B1* downstream of both *CCDC134* and *STT3A*. Additionally, the inability of CCDC134 overexpression to suppress the loss of STT3A (**Fig.3G**) suggests that CCDC134 depends on STT3A function to regulate HSP90B1 hyperglycosylation. These epistasis relationships allow the assembly of *CCDC134*, *HSP90B1* and *STT3A* into a provisional genetic pathway (**Fig.3J**) that can communicate the integrity of N-glycosylation to reception of WNT signals at the cell surface. This pathway model served as a framework for our subsequent biochemical studies.

### An N-terminal unstructured peptide in HSP90B1 regulates its own N-glycosylation

Inspection of the amino acid sequence around the facultative sequons in HSP90B1 failed to reveal any common sequence features. In fact, CCDC134 was able to suppress the N-glycosylation of ectopic, non- native sequons introduced into buried or exposed regions of HSP90B1, suggesting that the immediate sequence context around the sequons was not relevant (**fig.S4A**). So, what sequence features of HSP90B1 are required for its regulation by CCDC134? To simplify glycosylation analysis by gel shifts, we used a HSP90B1 variant carrying only three of the facultative sequons located in its middle (M) domain (**Fig.4A**). The remaining two facultative sequons (N62 and N107, which are not highly N- glycosylated in the absence of CCDC134 based on our mass spectrometry data shown in **Fig.3D**) and the constitutive sequon (N217) were removed by mutagenesis.

**Figure 4.**
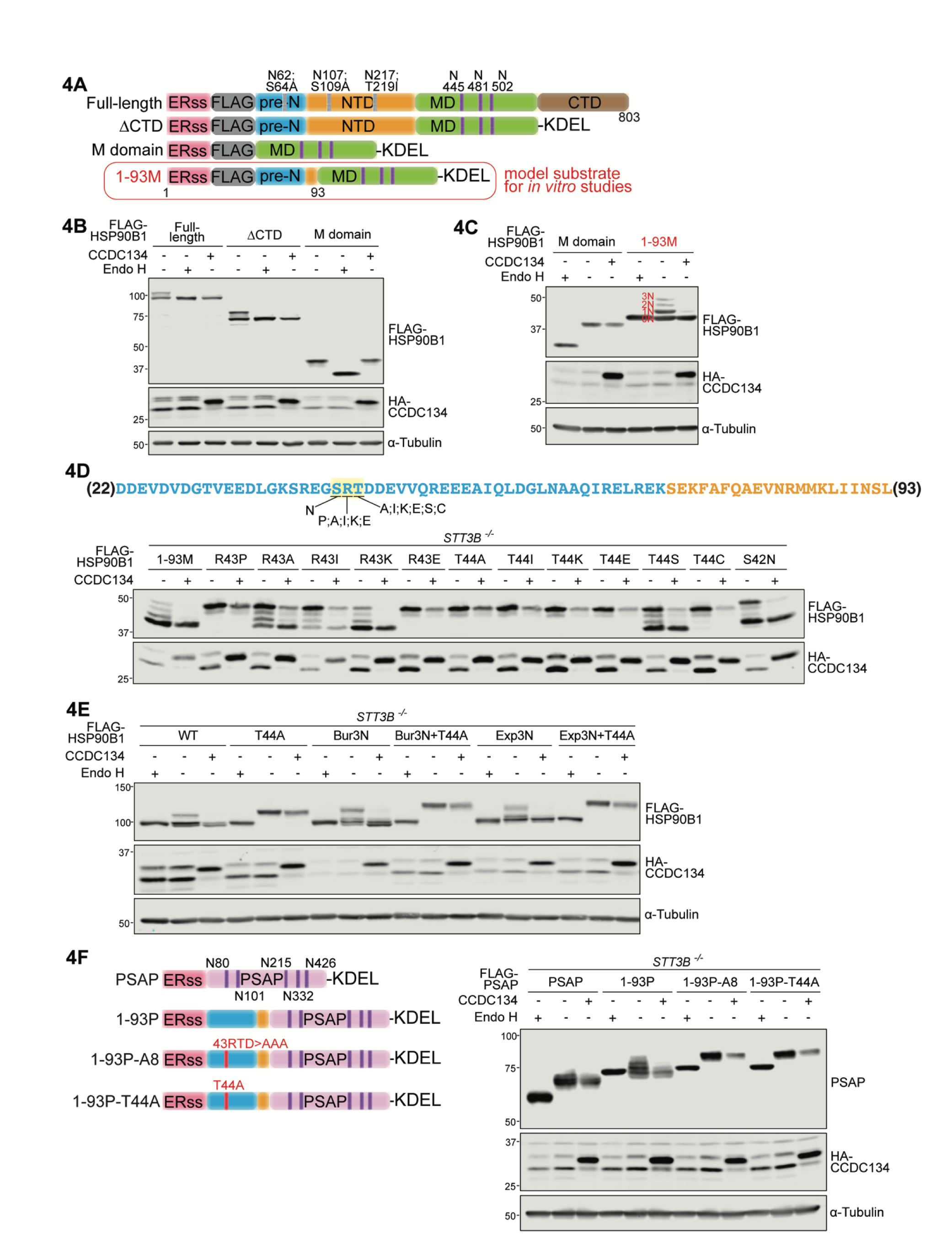
The pre-N segment of HSP90B1 regulates its own N-glycosylation. (**A**) Variants of HSP90B1 used for cell-based and *in vitro* assays in **Fig.4** and **5**, diagrammed following the scheme shown in **Fig.2E**. Key features include the ERss, ER signal sequence; FLAG, 3xFLAG tag; pre-N, unstructured segment; NTD, N-terminal domain; MD, middle domain; CTD, C-terminal domain. The N-glycosylation sites in pre-N and NTD were eliminated to allow easy assessment of the glycan modification of all three sequons in the M domain by gel shifts (see **C** and text). (**B**, **C**) Glycosylation status of HSP90B1 variants shown in (**A**) was assessed using gel shifts and Endo H sensitivity (see **Fig.2C**) after transient co-expression in HEK293T cells with WT CCDC134 (**+**) or a non-functional variant (-) lacking its ER signal sequence (see **Fig.1F**). The four predicted N-glycoforms (carrying 0, 1, 2 or 3 glycans) are labeled in (**C**) as 0N-3N in red lettering. (**D**) Glycosylation status of the 1-93M variant (see **A**) of HSP90B1 carrying the indicated mutations in the “SRT” motif (underlined in the sequence of the pre-N segment of HSP90B1 shown above). Each construct was co-expressed with functional (**+**) or non-functional (-) CCDC134. See **fig.S4** for deletion analysis and alanine scanning mutagenesis of the pre-N segment. (**E**) Glycosylation status of full-length (see **A**) FLAG-HSP90B1 carrying a T44A mutation in the “SRT” pseudosubstrate site identified in **D**. Bur3N and Exp3N refer to variant proteins containing three additional artificial sequons predicted to be buried (Bur) or exposed (Exp) based on the PDB 5ULS HSP90B1 structure(*58*). (**F**) Glycosylation status of chimeric proteins constructed by fusing the 1-93 segment of HSP90B1 (or mutants within the “SRT” pseudosubstrate site) to the obligate OST-A substrate PSAP, which contains five N-glycosylation sites shown in the domain diagram. T44A changes “SRT” to “SRA” and A8 changes “RTD” ro “AAA” (see **Fig.4D** and **fig.S4C**). All chimeras carry the ERss of HSP90B1. Note that in **D**, **E** and **F**, all constructs were transiently expressed in *STT3B^-/-^* HEK293T cells to exclude any contribution from OST-B glycosylation. See also **fig.S4**.

The isolated M domain of HSP90B1 was efficiently glycosylated (**Fig.4B**). CCDC134 co-expression had no effect on M domain glycosylation, suggesting that regions of the protein distant from the sequons themselves were required for preventing their glycosylation (**Fig.4B**). The C-terminal domain (CTD) was not required for CCDC134 regulation, thereby implicating the N-terminal domain (NTD) and the pre-N peptide segment. Systematic deletion analysis showed that amino acids 1-93 of HSP90B1, which includes the ER signal sequence and all of the pre-N segment, could affect the N-glycosylation of distant sequons in the M domain (**fig.S4B**). Deletions that extended into this 1-93 segment led to two distinct consequences: they increased the extent of M domain hyperglycosylation and abolished the capacity of CCDC134 to suppress this hyperglycosylation. Interestingly, deletions in the pre-N segment have previously been noted to affect HSP90B1 glycosylation and chaperone activity in cells (*32*, *58*). A minimal model substrate, in which the unstructured N-terminal 93 amino acids of HSP90B1 are fused to its M domain carrying three facultative sequons, was used for many of our subsequent experiments (hereafter called 1-93M, **Fig.4A**). The hyperglycosylation of 1-93M was efficiently suppressed by CCDC134 and its shorter length enabled efficient translation and glycosylation analysis by gel electrophoresis (**Fig.4C**).

To identify specific amino acid residues within the pre-N sequence that mediate regulation by CCDC134, we conducted alanine scanning mutagenesis. Sequential sets of three residues from amino acids 22 to 93 (excluding the signal sequence) in 1-93M were mutated to alanine, and residues that were originally alanine mutated to serine. Each of these twenty four mutants (A1-A24) were then evaluated for (1) the extent of N-glycosylation and (2) whether their glycosylation could be suppressed by CCDC134 co-expression (**fig.S4C**). Triplet alanine mutations in an 18 amino acid stretch of the pre-N domain close to the signal sequence markedly increased the efficiency of glycosylation, resulting in 1-93M that was fully glycosylated at all three sites in the M domain (**fig.S4C**). In contrast, wild-type 1-93M was a mixture of species carrying between 0-3 glycans, with the predominant form being unglycosylated. In addition, mutations that enhanced N-glycosylation of 1-93M were also much less sensitive to CCDC134 co-expression (**fig.S4C**).

The unexpected observation that a peptide segment can regulate the N-glycosylation of distant sequons that follow it suggested a substrate-directed auto-inhibitory model: the N-terminal 1-93 sequence, which emerges from the SEC61 translocon into the ER lumen before the M domain, inhibits the ability of OST-A to glycosylate any subsequent sequons in the same polypeptide. Inspection of the sequence where triplet mutations markedly enhance N-glycosylation (and prevent CCDC134 regulation) revealed a potential pseudosubstrate sequon (serine-arginine-threonine or “SRT” instead of asparagine- arginine-threonine or “NRT”) that might bind and inhibit the catalytic activity of STT3A (**Fig.4D**). This model is consistent with the observations that mutations in the sequon binding site in STT3A, predicted to be important for engaging a pseudosubstrate inhibitor, are impaired in their ability to suppress HSP90B1 hyperglycosylation (**Fig.3H**). Within the “SRT” motif, mutation of the R to proline (P) abolished activity (**Fig.4D**), mirroring the known fact that P in the middle position of classical sequons are not tolerated (*59*). However, the charge reversal of R to glutamic acid (E) also abolished activity (**Fig.4D**), showing that the pseudosubstrate sequence has more stringent requirements than a native sequon.

Mutation of the third T to serine (S) is well-tolerated, but its alteration to several other residues abolished activity (**Fig.4D**), again mirroring the sequence rules for classical sequons. Strikingly, a single conservative change of the T in SRT to alanine (hereafter “T44A”) resulted in near-complete N-glycosylation of all three sequons in the M domain of 1-93M and also prevented their regulation by CCDC134 (**Fig.4D**). The T44A mutation also abolished the ability of CCDC134 to suppress the N-glycosylation of full-length HSP90B1, both at native facultative sequons and at ectopically introduced artificial sequons (**Fig.4E**). Notably, the T44A point mutation also dramatically increased the extent of N-glycosylation in the context of full-length HSP90B1, leading to complete glycan occupancy (**Fig.4E**).

Interestingly, when wild-type HSP90B1 is expressed in *STT3B*^-/-^ cells, we see two distinct bands that correspond to the fully unglycosylated and monoglycosylated forms (**Fig.4E**), suggesting that the N217 constitutive sequon can also be glycosylated post-translationally by OST-B (*30*), consistent with its presence on a surface-exposed loop (**Fig.2F**).

As a stringent test of the model that the N-terminal 1-93 peptide segment of HSP90B1 functions as a pseudosubstrate inhibitor of the OST-A complex, we transplanted this sequence to PSAP, a protein that carries five sequons glycosylated exclusively by OST-A (**Fig.4F**). As expected, wild-type PSAP was efficiently N-glycosylated and insensitive to CCDC134 co-expression. A chimeric PSAP with the 1-93 HSP90B1 segment encoded at the N-terminus (1-93P) was glycosylated much less efficiently than wild-type PSAP, as seen by a smaller gel shift, consistent with the model that the 1-93 sequence from HSP90B1 can partially suppress the glycosylation of downstream sequons. CCDC134 co-expression was able to further suppress N-glycosylation of 1-93P. The T44A point mutation in the pseudosubstrate site of 1-93P restored the ability of OST-A to fully N-glycosylate PSAP and also prevented the suppressive effect of CCDC134 (**Fig.4F**), similar to the effect of this mutation on the 1-93M model substrate (**Fig.4D**) or on full-length HSP90B1 (**Fig.4E**).

We conclude that the N-terminal ∼93 amino acids of HSP90B1, predicted to be largely unstructured, can inhibit the N-glycosylation of the facultative sequons that follow it, perhaps by functioning as a pseudosubstrate inhibitor of STT3A. Notably, this effect is seen even in the absence of CCDC134: this unstructured peptide can partially suppress N-glycosylation when fused to the M domain of HSP90B1 or when fused to an entirely different protein (**Fig.4C,4F**). This same sequence is also required for the ability of CCDC134 to completely abolish N-glycosylation. These results led us to consider the hypothesis that the hyperglycosylation of HSP90B1 is regulated by a complex between its own N-terminus, CCDC134 and OST-A.

### Co-translational recruitment of CCDC134 to the ER translocon by the HSP90B1 nascent chain

To investigate biochemical interactions between CCDC134, HSP90B1 and OST-A, we used an established cell-free translation and translocation system that combines rabbit reticulocyte lysate (RRL) with microsomes isolated from HEK293T cells of any desired genotype (*60*). This system can be programmed with mRNA encoding a single protein, and both translation and translocation can be monitored in a time-resolved manner without interference from the multitude of mRNAs present in cells.

This system has been used extensively to study co-translational translocation of proteins into the ER and for the biochemical and structural characterization of the ER translocons, including the translocon formed by the association between SEC61 and OST-A (*12*, *61–63*).

Reactions containing wild-type, *CCDC134*^-/-^, *STT3A*^-/-^ or *STT3B*^-/-^ microsomes were programmed with mRNA encoding the 1-93M model substrate (**Fig.5A,5B**) derived from HSP90B1. As we observed in cells, N-glycosylation of 1-93M was enhanced when it was translated in the presence of microsomes lacking either CCDC134 or STT3A (**Fig.5B**). Furthermore, we could validate that the hyperglycosylation of 1-93M was indeed co-translational in the presence of *CCDC134*^-/-^ microsomes or when the T44A mutation was introduced in the putative pseudosubstrate site. Co-translational glycosylation was assessed by the Endo H sensitivity of 1-93M translation products that were shorter than the full-length protein (**fig.S5A,S5B**). We infer that these represent nascent peptides in the process of being translated and translocated into the ER. Radioactive pulse experiments in cells confirmed that hyperglycosylated HSP90B1 was detected at the earliest time points after the addition of ^35^S-Methionine, concomitant with the detection of monoglycosylated HSP90B1 (**fig.S5C**). Pulse-chase experiments failed to reveal any evidence of label transfer from monoglycosylated to hyperglycosylated HSP90B1 (**fig.S5D**). Both observations are consistent with co-translational N-glycosylation, considered the dominant mode of glycosylation by OST-A.

**Figure 5.**
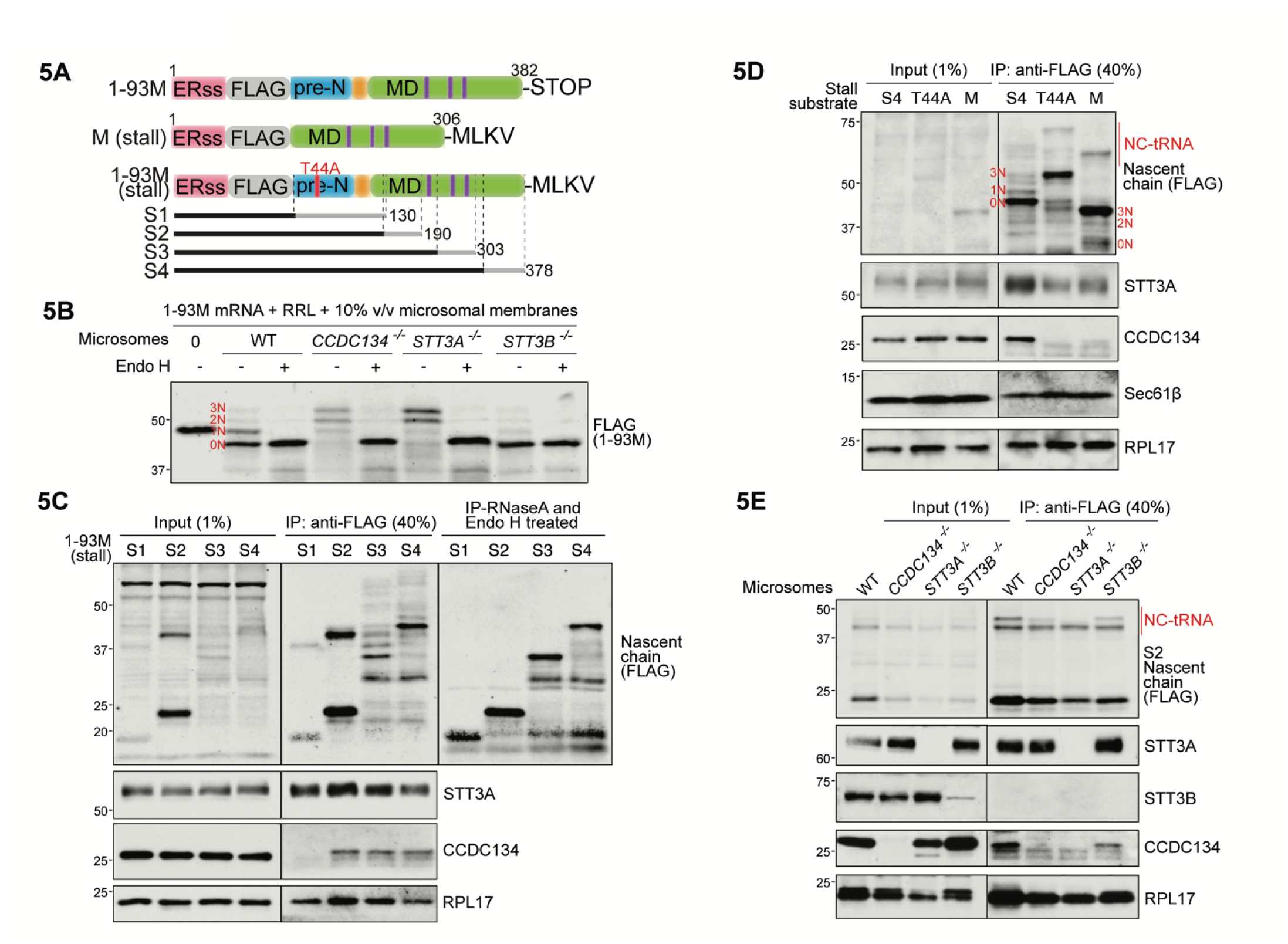
The pre-N segment of HSP90B1 recruits CCDC134 to the translocon to prevent N- glycosylation of distal sequons. (A) Constructs used for *in vitro* translation experiments. For the 1-93M (stall) constructs, the dark gray indicates the region predicted to be fully in the ER lumen and light gray indicates the region predicted to be in the translocon and ribosome exit tunnel. (B) Glycosylation status of the 1-93M variant of HSP90B1 (see **Fig.4A**) translated in rabbit reticulocyte lysate (RRL) in the presence of rough microsomal membranes generated from wild-type or clonally- derived *CCDC134^-/-^*, *STT3A^-/-^*, or *STT3B^-/-^*HEK293T cells and concentrated by immunoprecipitation on anti-FLAG beads. The four N-glycoforms of 1-93M are labeled (compare to **Fig.4C**) and show sensitivity to Endo H treatment. (C) Association of endogenous CCDC134 with truncation variants of 1-93M lacking a STOP codon translated in the presence of rough microsomal membranes generated from wild-type HEK293T cells. The stalled nascent chain was immunoprecipitated (IP) using anti-FLAG beads and association with STT3A, CCDC134, and the ribosome (RPL17) assessed by immunoblotting. To clearly visualize the sizes of the truncation variants, each sample was also treated with RNase A to release tRNA and Endo H to collapse glycoforms (top right immunoblot). (D) Association of endogenous CCDC134 with stalled 1-93M, M domain alone or a 1-93M variant carrying a T44A mutation in the “SRT” pseudosubstrate site, see **Fig.4D**), assessed as in **B**. NC-tRNA: nascent chain-tRNA conjugates. (E) Association of endogenous CCDC134 with a stalled 1-93M nascent chain translated in the presence of microsomes isolated from wild-type (WT), *CCDC134^-/-^*, *STT3A^-/-^*, or *STT3B^-/-^* HEK293T cells, assessed is in **B**.

To assess biochemical interactions, we used various FLAG-tagged 1-93M constructs lacking stop codons to stall translation, trapping the nascent chains in the ER translocon (**Fig.5A**). The stalled nascent chain was isolated using the FLAG tag and associated proteins detected by immunoblotting. A series of stalled truncation mutants of 1-93M co-precipitated both STT3A and ribosome subunits, indicating association with a secretory translocon (**Fig.5C**). However, CCDC134 was recruited only when the stalled nascent chain was long enough such that the 22-93 pre-N sequence would be predicted to be exposed to the ER lumen and hence available to interact with the sequon binding site of OST-A (**Fig.5C**). The recruitment of CCDC134 to the nascent chain-translocon complex depended on the integrity of the pseudosubstrate SRT site in the pre-N domain of HSP90B1 (**Fig.5D**) and on the presence of STT3A (**Fig.5E**). Introduction of the T44A mutation in the SRT site dramatically increased N-glycosylation and abolished the recruitment of CCDC134 (**Fig.5D**). Taken together, cell-free reconstitution demonstrated that CCDC134 was recruited to the ribosome-translocon complex in a manner that was dependent on both OST-A and the pre-N segment of the HSP90B1 nascent chain.

Evolutionary sequence analysis provides support for a shared function of CCDC134 and the pre- N segment of HSP90B1 (**Fig.6A and fig.S6**). HSP90B1 is a member of the HSP90 family of dimeric ATP-dependent chaperones that are conserved across the tree of life (**fig.S6**). Within the HSP90 family, HSP90B1 belongs to the endoplasmin clade characterized by three unique sequence features: a signal sequence for targeting to the ER, a KDEL or equivalent ER-retention sequence, and the unstructured pre-N segment that is a focus of this work (**fig.S6**). While endoplasmins date back to the base of the eukaryotic tree, CCDC134 is a much later invention, found in animals and their sister groups, the choanoflagellates and filastereans, and the thecamonads, a group of eukaryotes related to both animals and fungi (**Fig.6A**). It is only within the CCDC134-containing group of organisms that we observe significant constraints on the sequence of the HSP90B1 pre-N domain, including the pseudosubstrate site and surrounding sequence (**Fig.6A**). Interestingly, fungi show a corresponding loss of both CCDC134 and sequence conservation in the pre-N segment (**Fig.6A** and **fig.S6**).

**Figure 6.**
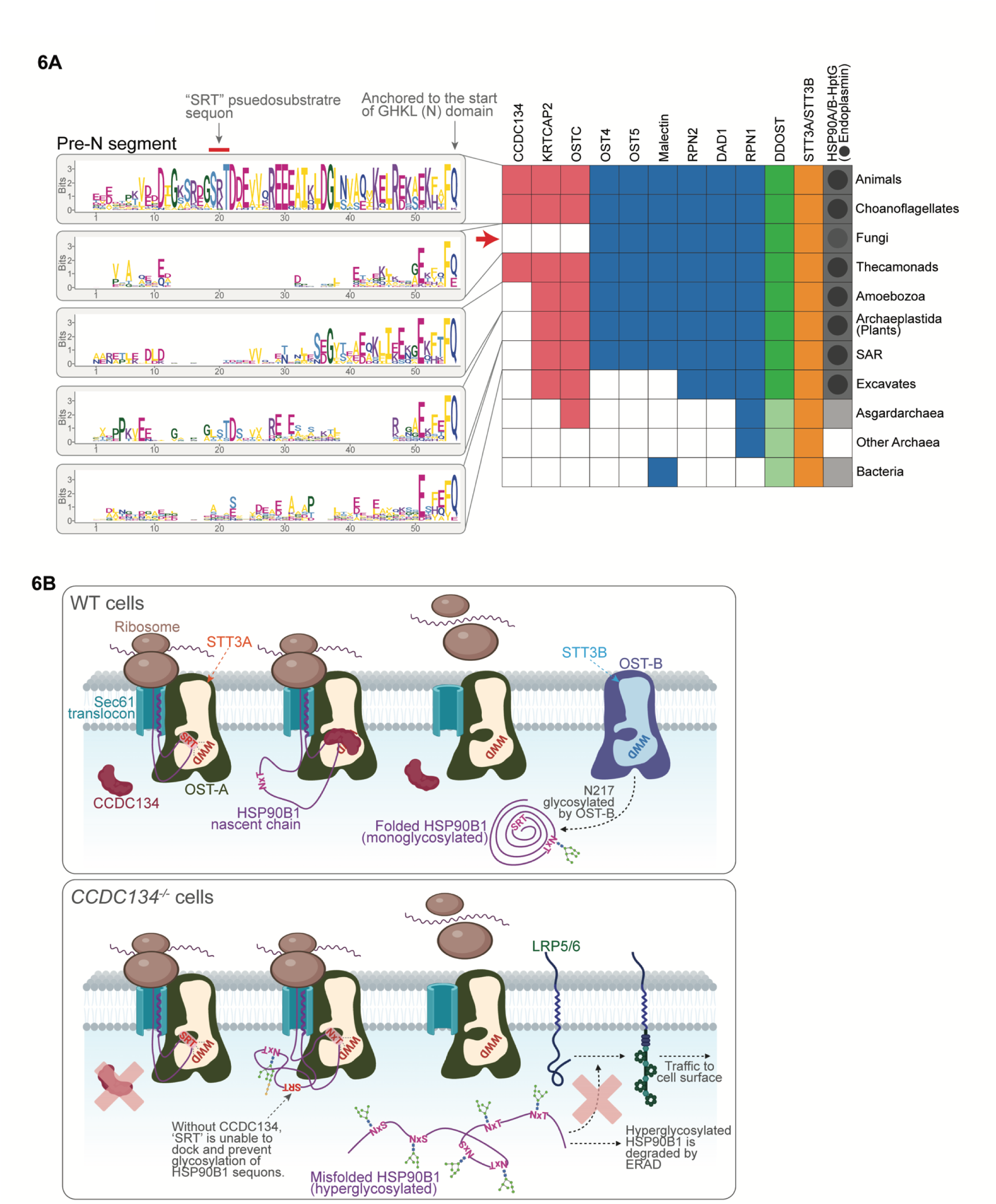
Evolutionary analysis of the pre-N segment of HSP90B1 and molecular mechanism of CCDC134 function. **(A)** Conservation of CCDC134, non-catalytic OST subunits, STT3A/B, HSP90 and endoplasmin (which includes HSP90B1 in mammals) across evolution. Shading indicates the presence of the protein in a specific sub-clade, and a black dot in the HSP90 column indicates the presence of a member of the endoplasmins. The conservation pattern of the pre-N segment in each sub-clade is shown as a sequence logo to the left. The logos are anchored to the start of the HSP90 NTD (the ATP-binding GHKL domain) and the residues are scaled according to the conservation bit score. The concomitant loss of CCDC134, STT3A-specific subunits OSTC and KRTCAP2 that anchor OST-A to the SEC61 translocon, and HSP90B1 in fungi is highlighted with a red arrow. The complete HSP90 phylogenetic tree is shown in **fig.S6**. **(B)** A model for the regulation of HSP90B1 hyperglycosylation (see text for details).

## DISCUSSION

The ribosome-translocon complex, originally thought to serve as a static conduit for transport of a peptide chain across the ER membrane, is emerging as an unexpectedly dynamic assembly of factors that control the fate of secreted and membrane proteins (*64*). Recent studies show that the compositions of the translocons that assemble to promote the insertion of secreted proteins with signal peptides, single-pass transmembrane proteins and multi-pass membrane proteins can be very different (*61*, *62*). Cryo-electron tomography (Cryo-ET) provides direct *in situ* evidence for polysomes docked to translocons composed of different components, with the most abundant containing the SEC61 protein channel, the TRAP complex and OST-A (*65*). In these cases, properties of the nascent chain substrate itself seems to direct the assembly of translocon components required for biogenesis. Our work is consistent with this view that custom translocons, tailored by information provided by the nascent chain, can serve as regulatory platforms. In our case, the nascent chain recruits CCDC134 to an OST-A-containing translocon to regulate the N-glycosylation and stability of the ER chaperone HSP90B1, with direct consequences for WNT signaling sensitivity and tissue development (**Fig.6B**).

We speculate that recruitment of other factors like CCDC134 to OST-A provide an unexplored mechanism by which translocon composition can control the biogenesis, itinerary and function of substrates. This possibility is consistent with the evolutionary history of the N-glycosylation apparatus. While the glycosyltransferase catalytic subunits (STT3A, STT3B) and DDOST can be traced back to the last universal common ancestor, the other subunits were added later in evolution (**Fig.6A**). While some of these distinguished the archaeo-eukaryotic complexes from the bacterial versions (e.g., RPN1, OSTC), the origin of eukaryotes saw an increase in complexity with multiple additional subunits (e.g., KRTCAP2). The emergence of CCDC134 later in evolution indicates that this process of complexification continued within eukaryotes, thereby adding a further layer of regulation to N-glycosylation.

An exciting prospect raised by these observations is that translocon composition and function may be regulated by signals from the ER or the cytoplasm to regulate cell-cell communication in multicellular organisms. HSP90B1 is not required for the growth or survival of individual cells in culture, but it is essential for the survival of all metazoan embryos in which its requirement has been tested (*66*). Mouse ES cells lacking HSP90B1 are viable and can differentiate into all three germ layers; however, mouse embryos lacking HSP90B1 die at gestational day 7 before gastrulation (*67*). These observations suggest that HSP90B1 is important for proper cell-cell communication during tissue development or homeostasis, but is dispensable for cell autonomous processes. Indeed, the known client list of HSP90B1 is enriched in cell surface receptors and secreted ligands. Apart from LRP5/6, HSP90B1 assists the folding and ER export of a subset of immunoglobulins, Toll-like receptors, integrins, insulin-like growth factors, the TGF-β docking receptor GARP and the platelet glycoprotein Ib complex (*3*, *66*). Our work shows that the function of HSP90B1 itself can be controlled by N-glycosylation. Based on its link to the human disease OI, this regulatory mechanism may be particularly important in professional secretory cells like osteoblasts, which face high demands on the N-glycosylation machinery.

Differentiation along the osteoblast lineage is characterized by massive expansion in the capacity of the ER to synthesize, N-glycosylate and secrete components of the bone matrix.

Our work also reveals an unexpected divergence in the function of the OST-A and OST-B complexes, previously thought to function together to maximize N-glycosylation of secretory pathway proteins. In our case, OST-A and OST-B work in opposition: OST-B promotes hyperglycosylation and degradation of HSP90B1 while OST-A serves a protective role. We propose that the pre-N segment tethers HSP90B1 to translocon-associated OST-A, protecting it from OST-B until it completes folding and renders its facultative sequons inaccessible (**Fig.6B**). This tethering is mediated through an interaction between the pre-N segment, CCDC134 and the sequon-binding site of STT3A. While the sequon-binding site is also conserved in STT3B, this model critically depends on the N- to C-terminal scanning mechanism of co-translational glycosylation, a feature unique to OST-A. Our model explains several perplexing observations: (1) catalytically inactive OST-A can still suppress HSP90B1 hyperglycosylation, suggesting a scaffolding rather than enzymatic function (**Fig.3H**), (2) the requirement for OSTC (**Fig.3B**), which tethers OST-A to the translocon but is not required for its integrity or oligosaccharyltransferase activity (*10*), and (3) mutations in the sequon binding site, but not the LLO binding site or active site, in STT3A allow HSP90B1 hyperglycosylation by OST-B (**Fig.3H, 3I**). Intriguingly, a non-catalytic scaffolding role for both OST-A and OST-B has also been proposed during dengue virus infection (*68*). It is possible that the association of OST-A to the translocon, through the dedicated subunits OSTC and KRTCAP2, enabled an expansion of its functions (beyond N-glycosylation) to include serving as a scaffold to recruit ER factors to proteins being co-translationally transported across the ER membrane. Consistent with this model, recent Cryo-ET data reveals presence of co-factors associated with OST-A (*65*).

### Limitations of this study

While we focused on WNT signaling, it is likely that HSP90B1 hyperglycosylation and degradation in the absence of CCDC134 affects multiple clients of HSP90B1. This possibility is reinforced by the observation that CCDC134 is present in choanoflagellates, filastereans, and thecamonads, lineages that lack WNT signaling (**fig.S6**). Thus, we expect that the loss of CCDC134 may impact other cell-cell communication pathways involved in development (e.g. IGF-1 signaling (*67*)) or immune function (*3*). For example, *CCDC134* was identified as a gene regulating T helper 2 (TH2) differentiation (*69*), though the mechanism remains unknown. A full understanding of the biological functions of CCDC134 will likely require its selective disruption in specific tissues, cell lineages and temporal contexts.

Future research should also focus on how CCDC134 is itself regulated. We have found that CCDC134 mRNA or protein levels are not regulated by either WNT or unfolded protein response pathways, suggesting that it is not a feedback regulator of WNT signaling or directly responding to ER stress (though it is able to attenuate ER stress induced HSP90B1 hyperglycosylation when overexpressed). Public databases do not suggest that CCDC134 expression is highly restricted to specific tissues, but further investigation with CCDC134-specific probes is needed. We anticipate that regulation of CCDC134 (by expression, ER targeting or post-translational modification) can be used to control the sensitivity of a cell to WNT ligands (and thus to control proliferation and differentiation outcomes).

We end by speculating that the N-glycosylation of other secretory pathway proteins may be regulated by pre-N-like peptide segments within their sequences, perhaps in cooperation with CCDC134- like ER adapters, representing a new mechanism of receptor regulation and signal transduction.

## ACKNOWLEDGEMENTS

RR was supported by grants from the National Institutes of Health (GM118082, HD101980), MM by an American Cancer Society – Jean Perkins Foundation Postdoctoral Fellowship in Cancer Research, PF- 20-121-01-TBE, A.P. Giannini Foundation Postdoctoral Fellowship, and Stanford School of Medicine Dean’s Postdoctoral Fellowship. We are grateful for support from Grant P41 GM108538 (National Center for Quantitative Biology of Complex Systems) to JJC and the French association of osteogenesis imperfecta (AOI) and the Fondation Philanthropia to VC-D.

## AUTHOR CONTRIBUTIONS

MM and RR conceived the project. MM and RR wrote the manuscript. Experiments related to CRISPR/Cas9 screens and validation were performed by MM, RD, GVP. Analysis of CRISPR/Cas9 screen results was performed by RD. Patient OI fibroblasts were provided by VCD. Mass spectrometry experiments were performed and analyzed by AJ with technical assistance from ES, KAO, and guidance from JJC. Evolutionary analyses were performed by LA. All other experiments were performed by MM with guidance from RR.

## COMPETING INTERESTS

The authors declare no competing interests.

## DATA AND MATERIALS AVAILABILITY

All unique reagents generated in this study are available from the Lead Contact, Rajat Rohatgi, and will be provided upon request. All data are available in the manuscript or the supplementary materials. This paper does not report original code. All FASTQ files from CRISPR/Cas9 screen have been deposited into the NIH Short Read Archive (SRA) with BioProject accession number PRJNA1087438. Raw data files for N-glycoproteomic analyses have been deposited into the MassIVE database under the accession number MSV000094280. All patient fibroblast cells used in this manuscript have been previously established by Dr. Valérie Cormier-Daire and reported in a prior peer-reviewed publication from her laboratory (PMID 32181939, J Bone Miner Res. 35(8) 1470-1480). As noted in this original publication, informed consent for participation and sample collection were obtained via protocols approved by the Necker-Enfants Malades Hospital, AP-HP, F-75015 Paris, France. This work used only these previously published unidentifiable or de-identified cell lines and did not involve any active patient recruitment or new sample collection from patients.

## SUPPLEMENTARY MATERIALS

Materials and Methods

Figs. S1 to S6

References (*71–84*)

Data S1 to S2

**The PDF file includes:**

Materials and Methods

Figs. S1 to S6 References (*71–84*)

**Other Supplementary Materials for this manuscript include the following:**

**Data S1**. Results from genome-wide CRISPR/Cas9 knockout screens.

**Data S2**. Results from N-glycoproteomics.

## MATERIALS AND METHODS

### Constructs

*CCDC134* constructs: *CCDC134* cDNA with a 3xHA tag inserted just after the ER signal sequence was ordered as a gene fragment from Twist Bioscience, cloned into the Gateway compatible entry vector pENTR2B Dual Selection Vector, and used as a template for generation of all *CCDC134* constructs.

CCDC134 constructs were cloned into pEF5/FRT/V5-DEST (Thermo Fisher Scientific) or pLenti CMV PURO DEST (*71*) using Gateway recombination methods. Doxycycline inducible *CCDC134* constructs were cloned by PCR amplification followed by Gibson assembly (New England Biolabs) into pLenti-TRE- rtta3G-BLAST.

*HSP90B1* constructs: pDONR223_HSP90B1_WT was purchased from Addgene (Plasmid #82130) and Gibson assembly was used to insert a 3xFLAG tag just after the ER signal sequence. All HSP90B1 mutant and truncation constructs were generated using PCR amplification followed by Gibson assembly and cloned into pEF5/FRT/V5-DEST (Thermo Fisher Scientific, Invitrogen) or pLenti CMV PURO DEST (*71*) using Gateway methods. The following mutations were made in HSP90B1: HSP90B1^1N^ (T219I), HSP90B1^5N^ (S64A, S109A, S447A, T483I, T504I), HSP90B1 with ectopic buried (Bur3N) sequons (G164S, K493T, E659T), and HSP90B1 with ectopic exposed (Exp3N) sequons (L145T, M622T, Y678T). *STT3A* constructs: STT3A-FLAG was a gift from Jan Carette (*68*) and used as a template to generate STT3A constructs and mutants.

All constructs were fully sequenced to confirm accuracy.

### Reagents and antibodies

WNT3A conditioned media was produced as previously described (*20*). Briefly, 1.5 x 10^6^ Wnt-3A cells or L cells were seeded in 15 cm tissue culture-treated dishes, grown for 3 days, media was replenished, and after an additional 3 days, conditioned media was collected, filtered through 0.2 µm polyethersulfone (PES) membrane filter, aliquoted, flash-frozen in liquid nitrogen, and stored at −80°C. Puromycin and blasticidin were purchased from Sigma-Aldrich. The transfection reagent polyethylenimine (PEI) was purchased from Polysciences and polybrene from MilliporeSigma. Bafilomycin A1 was purchased from Cayman Chemical, Bortezomib from LC labs, kifunensine from Tocris Bioscience, and NMS-873 from Sigma-Aldrich. Endoglycosidase H (EndoH) and Peptide-*N*-Glycosidase F (PNGaseF) were purchased from New England Biolabs. Phenylmethanesulfonyl fluoride (PMSF) and cycloheximide were purchased from Sigma-Aldrich, micrococcal nuclease from New England Biolabs, RNase-free DNase from Promega, SUPERaseIn from Invitrogen, and RNaseA from Gold Biotechnology. ANTI-FLAG® M2 Affinity Gel was purchased from Sigma-Aldrich. The following primary antibodies were used: mouse anti- CCDC134 (E-5, Santa Cruz Biotechnology, 1:500); mouse anti-HSP90B1 (H-10, Santa Cruz Biotechnology, 1:2000); mouse anti-LRP5 (B-9, Santa Cruz Biotechnology, 1:500); rabbit anti-LRP6 (C5C7, Cell Signaling Technology, 1:1000 for immunoblot); mouse anti-LRP6 ectodomain (clone A59, MilliporeSigma, 2ug/sample for cell surface staining); Alexa Fluor® 488 rabbit anti-Giantin (A488-114L, Covance, 1:500 for immunofluorescence); mouse anti-ɑ-Tubulin (Clone DM1A, MilliporeSigma, 1:10000); rabbit anti-active (non-phosphorylated) β-catenin (D13A1, Cell Signaling Technology, 1:500); rabbit anti-Na/K ATPase (3010S, Cell Signaling Technology, 1:1000); rabbit anti-MESD (10958-1-AP, Proteintech, 1:1000); mouse anti-FLAG (clone M2, MilliporeSigma, 1:2000); rabbit anti-FLAG (F7425, Sigma-Aldrich, 1:2000); rabbit anti-PSAP (GTX101064, GeneTex, 1:1000); rabbit anti-STT3A (12034-1-AP, Proteintech, 1:1000); rabbit anti-STT3B (15323-1-AP, Proteintech, 1:1000); rabbit anti-OSTC (PA5-34060, Invitrogen, 1:1000); mouse anti-RPL17 (C-8, Santa Cruz Biotechnology, 1:2000); rabbit anti-Sec61b (15087-1-AP, Proteintech, 1:1000). Secondary antibodies conjugated to horseradish peroxidase or Alexa Fluor dyes were obtained from Jackson Laboratories and Thermo Fisher Scientific. IRDye® 800CW Donkey anti-mouse and anti-rabbit IgG (H + L) were purchased from LI-COR.

### Cell lines

L Wnt-3A (ATCC Cat. # CRL-2647), L cells (ATCC Cat. # CRL-2648), HEK293T (ATCC Cat. # CRL-3216), RKO cells (ATCC Cat. # CRL-2577), and C3H10T1/2 cells (ATCC Cat. # CCL-226) were grown in Dulbecco’s Modified Eagle Medium (DMEM) containing high glucose (Cytiva) and supplemented with 10% fetal bovine serum (FBS) (Sigma-Aldrich), 1 mM sodium pyruvate (Gibco), 2 mM L-Glutamine (Gibco), 1x MEM non-essential amino acids solution (Gibco), penicillin (40 U/ml) and streptomycin (40 μg/ml) (Gibco), in a humidified atmosphere containing 5% CO_2_ at 37°C. MC3T3-E1 cells (ATCC Cat. # CRL-2593) were grown in Alpha Minimum Essential Medium (aMEM) without ascorbic acid (Gibco Cat. # A1049001) supplemented with 10% FBS (Sigma-Aldrich), penicillin (40 U/ml) and streptomycin (40 μg/ml) (Gibco), in a humidified atmosphere containing 5% CO_2_ at 37°C. Clonally derived *STT3A^-/-^*and *STT3B^-/-^* HAP1 cells were a gift from Jan Carette(*68*) and the wild-type HAP1 cell line from which pooled knockout lines were generated was a gift from Thijn Brummelkamp (now available from Horizon Discovery, Cambridge, United Kingdom). All HAP1 cells and derivatives thereof were grown in Iscove’s Modified Dulbecco’s Medium (IMDM) (Cytiva Cat. # SH30228FS) and supplemented with 10% FBS (Sigma-Aldrich), 2 mM L-Glutamine (Gibco), penicillin (40 U/ml) and streptomycin (40 μg/ml) (Gibco), in a humidified atmosphere containing 5% CO_2_ at 37°C. Primary human fibroblasts were provided by Valerie Cormier-Daire and cultured as described previously(*5*).

Parent cell lines purchased from ATCC or Thermo Fisher Scientific (see above) came with a certificate of authentication from the vendor and were used without further validation. Patient fibroblasts from Dr. Cormier-Daire were validated by western blotting to ensure lack of CCDC134. HAP1 cells were validated by propidium iodide staining to ensure a haploid genome content. All stable or gene-edited cell lines derived from these parental cells were validated by western blotting or genomic PCR. Cell lines were confirmed to be negative for Mycoplasma infection when introduced into the lab (with the exception of patient fibroblasts).

### Pooled genome-wide CRISPR/Cas9 screens

*Generating a reporter cell line.* A clonally derived RKO WNT EGFP reporter (RKO-7TG_scc-12) was generated as described previously (*20*). The clonal reporter line was transduced with Cas9 (lentiCas9- Blast; Addgene#52962) and a second round of clonal derivation was performed to generate multiple clonal cell lines that were analyzed for optimal Cas9 expression and WNT3A-induced EGFP reporter fluorescence. A clonal cell line (RKO-7TG_scc-12; Cas9_scc-7) with >90% on-target genome editing activity using two positive control sgRNAs (*CTNNB1* and *TCF7L2*) and also displayed the widest dynamic range for WNT3A-induced EGFP reporter fluorescence was selected as the reporter line for CRISPR/Cas9 knockout screens. Screen design was modeled after previous screens in the lab (*72*). *Generating knockout library and screen.* The Brunello CRISPR library (Addgene #73178 (*73*)) was used to generate our genome-wide collection of mutant RKO cells. Brunello library amplification, lentiviral production, functional titer determination, and transduction were performed as described previously with minor modifications(*74*). Briefly, the Brunello library was amplified in Endura electrocompetent cells (Lucigen) and subjected to Next-Generation Sequencing (NGS) to determine sgRNA distribution. For lentivirus production, 18 million 293FT cells were seeded in T225 flasks (40 flasks in total) and transfected the following day with 3.4 μg pMD2.G (Addgene #12259), 6.8 μg psPAX2 (Addgene#12260), and 13.6 μg lentiviral target (CRISPR) plasmid, and 195 μl of 1 mg/ml polyethylenimine (Polysciences) per flask. 48 hours after transfection, lentivirus was harvested, filtered through a 0.45 μm filter, aliquoted into multiple 50 ml tubes and stored at -80°C. The functional titer of the lentivirus was determined by surviving RKO cells after 24 hours of infection with virus followed by 48 hours of 2ug/mL puromycin treatment. RKO-Wnt reporter cell line stably expressing Cas9 was transduced with the Brunello library at a Multiplicity of Infection (MOI) =0.3 (360 million cells were transduced with virus to achieve ∼1000 fold representation of each sgRNA) in the presence of 10 μg/ml polybrene. 24 hours after infection, cells were split and selected with puromycin (2 μg/ml) for seven days and frozen in aliquots of 5 million cells/vial. Genomic DNA (gDNA) was extracted from cells using Quick-gDNA Midiprep kit (Zymo Research) and subjected to NGS to determine sgRNA distribution. In all screens, 100 million cells were initially thawed into 5 x15 cm tissue culture-treated dishes and two days later split into a new set of 15 cm dishes such that at least 100 million cells were plated again to maintain 1000 fold representation of each sgRNA. On the fourth day the cells were split and 13 million cells were plated in 6 x 15 cm dishes. The next day the cells were treated with 50% WNT3A conditioned media for 24 hours. Cells were trypsinized and 4 million cells were pelleted and frozen (unsorted population) and the remaining ∼40 million cells (corresponding to 500-fold representation of each sgRNA in the Brunello library) were sorted for cells with the lowest 10% of EGFP fluorescence. The screen was performed twice under identical conditions.

*NGS sequencing and analysis.* Genomic DNA (gDNA) was extracted from unsorted and sorted cells and the sgRNA library was amplified as described previously (*74*). Briefly, each sgRNA library PCR was set up with a mix of 10 NGS-Lib-Fwd primers (“staggered” Fwd primer mix to increase library diversity) and a unique NGS-Lib-Rev primer with the entire gDNA (5 µg per 100 µL reaction). The PCR product was purified, quantified by qRT-PCR and subjected to sequencing on Illumina HiSeq. In all the screens, we averaged >100 reads per sgRNA in the library. For analysis, reads from the FASTQ files generated by sequencing were tallied for each guide by taking the first 20 base-pairs from each read (that were trimmed to remove adapter and vector backbone sequences) and mapping that sequence to the identical sgRNA sequence. For each screen, a table of reads per guide that includes the counts from the sorted and unsorted populations from both replicates was generated. The tables generated from the two independent duplicates of each screen were analyzed together by the MAGeCK computational tool (*75*), specifying the 1000 control sgRNAs for normalization and generation of the null distribution for MAGeCK with the “--control-sgrna” option and computing the log fold change for the gene using the mean of all of the guides for a given gene with the “--gene-lfc-method mean” option.

### Immunoblot analysis

For immunoblot analysis of β-catenin in RKO cells, whole cell lysates were prepared in β-catenin lysis buffer (*76*): 30 mM Tris at pH 7.4, 150 mM NaCl, 1% Triton X-100, 0.5 mM TCEP, 1 mM EDTA, 10% glycerol, 1mM NaF, 1 mM Na_3_VO_4_1, and 1x SIGMAFAST protease inhibitor cocktail (MilliporeSigma). For all other immunoblotting data presented in the manuscript, whole cell lysates were prepared in RIPA lysis buffer: 50 mM Tris at pH 8.0, 150 mM NaCl, 2% NP-40, 0.25% Deoxycholate, 0.1% SDS, 0.5 mM TCEP, 10% glycerol, 1x SIGMAFAST protease inhibitor cocktail (MilliporeSigma), and 1x PhosSTOP phosphatase inhibitor cocktail (Roche). For resolving LRP6 and full length HSP90B1 glycoforms, samples were run on a 7% Tris-glycine gel. For resolving 1-93M glycoforms, samples were run on a 9% Tris-glycine gel. The resolved proteins were transferred onto nitrocellulose membrane (Bio-Rad Laboratories) using a wet electroblotting system (Bio-Rad Laboratories) followed by immunoblotting.

### Flow cytometry analysis

For analysis of the WNT EGFP reporter, cells were treated with either WNT3A conditioned media or control conditioned media for 24 hours, trypsinized, filtered through 70 µm sterile cell strainer (Falcon), and EGFP fluorescence was measured immediately on a Sony SH800 flow cytometer. Fluorescence data for 10,000 singlet-gated cells was collected and subsampled thrice, each data point represents median reporter fluorescence from each subsampled population.

For cell surface staining of LRP6 in primary fibroblasts, cells were harvested by brief incubation with trypsin (3-4 min), immediately resuspended in complete media and counted. The cells were pelleted and resuspended in Staining Buffer or SB (10% FBS and 0.05% sodium azide prepared in PBS) at a concentration of 0.5 million cells/100 µL SB. 100 µL 10% donkey serum was added to 100 µL cell suspension and the cells were blocked for 10 min at room temperature. Cells were pelleted, resuspended in 100 µL fresh SB and stained with 2 µg of an anti-LRP6 antibody (A59, MilliporeSigma) for 30 min on ice. Cells were washed twice, resuspended in 100 µL fresh SB and then incubated for 30 min on ice in 1 ug of donkey anti-mouse IgG, Alexa Fluor 647 (Thermo Fisher Scientific). Cells were finally washed twice, resuspended in 100 µL fresh SB and analyzed on a Sony SH800 flow cytometer.

Fluorescence data for 2,000 singlet-gated cells was analyzed per experiment and the experiment was repeated thrice. For all other experiments, fluorescence data for 10,000 singlet-gated cells was collected unless indicated otherwise.

### Immunofluorescence analysis

RKO cells expressing doxycycline-inducible 3xHA tagged CCDC134 variants were seeded on coverslips, grown for 1 day, and treated with 25 nM doxycycline for 24 hours. Coverslips were washed with PBS, fixed with 4% (w/v) paraformaldehyde (PFA) at room temperature, permeabilized and blocked with PBS containing 1% donkey serum + 1% BSA + 0.1% Triton X-100 for 1 hour at 4°C, and incubated with primary antibodies (mouse anti-CCDC134 at 1:200 and rabbit anti-Giantin-AF488 at 1:500) for 1 hour at 4°C. Coverslips were washed and incubated with secondary antibody (donkey anti-mouse AF647), mounted in ProLong Diamond with DAPI (Invitrogen P36962), and images were collected using an Olympus IX83 epifluorescence microscope equipped with an Orca Fusion scMOS camera using a x100 oil objective (NA 1.45).

### Generation of clonally derived knockout cell lines

Clonal knockout RKO and HEK293T cell lines were generated using a dual sgRNA strategy as previously described (*72*). Briefly, two sgRNAs targeting candidate genes (or non-targeting control, NTC) 200-800 bases apart were designed using the Synthego Knockout Guide Design tool (http://design.synthego.com) and cloned into pSpCas9(BB)-2A-GFP (PX458; Addgene #48138) and pSpCas9(BB)-2A-mCherry, the latter generated by replacing the GFP cassette in PX458 with mCherry. Four days after co-nucleofection in RKO cells (Nucleofector 2b device using program A-024 and Lonza Cell Line Nucleofector® Kit V #VCA-1003) or co-transfection in HEK293T cells (X-tremeGENE9, Roche), GFP and mCherry double positive single cells were sorted into a 96-well plate using a Sony SH800 flow cytometer. Clonal lines were first screened by PCR to detect excision of the genomic DNA between the two sgRNA cut sites and further confirmed by immunoblot analysis using commercially available antibodies.

### Generation of CRISPR/Cas9-mediated pooled knockout cell lines

For validation of candidate genes from CRISPR/Cas9 screen, two independent guides were designed and individually cloned into lentiCRISPR v2 plasmid (Addgene #52961) (*77*). Lentivirus was produced as described above and used to infect RKO and HEK293T WNT-GFP reporter cells, followed by selection with puromycin (2 µg/mL) for 5 days. Pooled cell lines were analyzed by FACS for EGFP fluorescence after treatment with WNT3A or control conditioned media. For generation of pooled knockouts targeting all other genes in HAP1, MC3T3, and C3H10T1/2 cells, sgRNAs were selected from either the Brunello (human) or Brie (mouse) libraries and cloned into either lentiCRISPR v2 or lentiCRISPR v2-Blast (Addgene #83480). Lentivirus was produced and used to infect indicated cell lines, followed by selection with puromycin (2 µg/mL) or blasticidin (10 µg/mL).

### Generation of stable cell lines expressing transgenes

Stable addback cell lines expressing tagged CCDC134, tagged HSP90B1, or tagged STT3A were generated using the lentiviral expression system. To generate virus, 700,000 HEK293T cells were seeded onto a 6-well plate and 24 hours later transfected with 200 ng pMD2.G (Addgene), 400 ng psPAX2 (Addgene), and 800 ng of the desired pLenti CMV Puro DEST or pLenti-TRE-rtta3G-BLAST construct using 7 μl of 1mg/ml polyethylenimine (PEI) (Polysciences). Approximately 48 hours post transfection, lentivirus conditioned media was harvested and filtered through a 0.45 μm filter. 0.5 ml of the filtered lentivirus solution was mixed with 1.5 ml of complete media containing 8 μg/mL polybrene (MilliporeSigma). The diluted virus was then added to the indicated cells seeded on 6-well plates.

Approximately 24 hours post infection, cells were split and selected with puromycin (2 μg/mL) or blasticidin (10 µg/mL) for 3-7 days or until all the cells on the control plate are dead. For doxycycline inducible expression of 3xHA-CCDC134, cells were grown for 24 hours in a range of doxycycline concentrations with 5 nM inducing low, near-endogenous expression levels.

### Cell surface biotinylation assay

RKO cell lines were seeded at a density of 2 x 10^6^ on 10 cm tissue culture-treated dishes and grown for 2 days. Cell culture plates were removed from the 37°C incubator and placed on an ice-chilled metal rack in a 4°C cold room. Growth medium was removed and cells were quickly washed thrice with ice-cold DPBS+ buffer (1.47 mM KH_2_PO_4_, 8.06 mM Na_2_HPO_4_, 137.93 mM NaCl, 2.67 mM KCl, 0.9 mM CaCl_2_, 0.49 mM MgCl_2_.6H_2_O, 5.56 mM dextrose, and 0.33 mM sodium pyruvate). Cells were incubated with a freshly prepared solution of 0.4 mM Sulfo-NHS-SS-Biotin (Thermo Fisher Scientific) in DPBS+ buffer for 30 min. Unreacted Sulfo-NHS-SS-Biotin was quenched with Tris pH 7.4 at 50 mM for 10 min. Cells were then washed thrice with 1x Tris-buffered saline (25 mM Tris-HCl pH 7.4, 137 mM NaCl, and 2.7 mM KCl) and whole cell extracts were prepared in a buffer containing 50 mM Tris-HCl pH-7.4, 150 mM NaCl, 2% NP-40, 0.25% deoxycholate, 1x SIGMAFAST protease inhibitor cocktail (MilliporeSigma), and 1x PhosSTOP phosphatase inhibitor cocktail (Roche). Biotinylated proteins from clarified supernatants were captured on a streptavidin agarose resin (Solulink), washed, eluted in NuPAGE-LDS sample buffer containing 100 mM DTT at 42°C for 30 min to cleave and release biotinylated proteins, and assayed by immunoblotting.

### LC-MS/MS Proteomics

*Peptide Preparation.* Cell pellets were removed from -80 °C, where they were maintained prior to analysis. 1 mL of 5.4 M guanidine hydrochloride in 100 mM Tris HCl, pH 8.0, was added to each cell pellet. Samples were gently vortexed, then probe sonicated for 10 seconds to ensure cell pellet resuspension. A bicinchoninic acid (BCA) protein assay (Pierce, Rockford, IL) was performed according to manufacturer’s instructions to determine the protein concentrations. 1 mg of protein from each cell pellet was transferred into separate 1.5 mL microcentrifuge polypropylene tubes. Sufficient LC-MS grade methanol was added to each sample to bring each sample to 90% volume/volume methanol in composition, then vortexed for 10 seconds. Each sample was then centrifuged at 9,000 x g for 5 minutes at 5 °C to pellet the protein. After the supernatants were carefully decanted to waste. Each pellet was resuspended in freshly prepared 8M urea, 100 mM Tris HCl pH 8.0, 10 mM TCEP, 40 mM 2- chloroacetamide and vortexed for not less than 15 minutes at ambient temperature to resolubilize the protein. 20 µL of 1 mg/mL LysC prepared per manufacturer’s instruction (VWR, Radnor, PA) was added to each 1 mg protein sample, then allowed to incubate at ambient temperature for four hours while gently rocking. Samples were then diluted with freshly prepared 100 mM Tris HCl pH 8.0 to reach a final urea concentration of 2 M, after which 20 µL 1 mg/mL trypsin (Promega, Madison, WI) was added to each sample. Samples were incubated at ambient temperature overnight while gently rocking. To stop digestion, 20 µL TFA was added to each sample, after which samples were centrifuged at 9,000 x g for 5 minutes to pellet insoluble material. The resulting supernatant was desalted using Strata-X 33 µm polymeric reversed phase SPE cartridges (Phenomonex, Torrance, CA).

*Glycopeptide Enrichment.* The desalted peptides were dried down in a vacuum centrifuge (Thermo Fisher Scientific, Waltham, MA). For glycopeptide enrichment, 1 mg desalted peptides were resuspended in 90% acetonitrile in 1% TFA, then placed onto 10 mg SOLA SPE columns (Thermo Scientific). The peptides were enriched for N-glycopeptides according to the published protocol (*78*) without the use of the vacuum manifold. The resulting enriched mixture was dried down in a vacuum centrifuge and resuspended in 20 µL 0.2% formic acid in water for analysis.

*Instrument Analysis.* Sample analysis was performed using a Vanquish Neo HPLC system coupled to an Orbitrap Ascend Tribrid mass spectrometer (Thermo Scientific, San Jose, CA). Mobile phase A was water with 0.2% formic acid, and mobile phase B was 80:20 v/v ACN:H2O with 0.2% formic acid. The gradient elution was carried out with a flowrate at 0.300 µL/min. 1 µg of enriched peptides (N-glycoproteomics) or unenriched tryptic peptides (shotgun proteomics) were loaded onto a 75 µm i.d. column with 1.7 µm, 130 Å pore size, Bridged Ethylene Hybrid (BEH) C18 particles (Waters, Milford, MA), packed in-house to a length of 30 cm (*79*). The column was heated to 50 °C during analysis.

For N-glycoproteomics mass spectrometer analysis, positive mode ionization was used. MS1 scans were acquired from 0 to 90 minutes every second at a scan range of 300-2,000 m/z, with a resolution of 60,000 in the Orbitrap and maximum injection time of 123 ms; normalized AGC target (%) was set to 250, equivalent to 1e6 ions, and RF lens (%) set to 30. Precursor ions with charge states 2-6 were isolated from a 1.3 Da window in the quadrupole with a dynamic exclusion period of 20 seconds. Data-dependence HCD MS2 scans with 36% normalized collision energy, maximum injection time of 59 ms, and a normalized AGC target of 200% (equivalent to 1e5 ions) were acquired in the Orbitrap at a resolution of 30,000 for precursors with the defined first mass of 150 m/z, and scanned for trigger ions of 204.0867, 138.0545, 366.1396, 274.0921, 292.1027, 126.055, 144.0655, 168.0654 or 186.076 (± 10 ppm). If trigger ions were detected, the precursor was re-isolated and fragmented with sceHCD of 35±15 over the scan range of 150-4,000 with the resolution of 30k and max injection time of 100 ms(*80*).

For shotgun proteomics mass spectrometer analysis, with positive mode ionization, MS1 scans were collected every second in the Orbitrap with a scan range of 300-1,350 m/z from 0 to 90 minutes, with a resolution of 240,000, a maximum injection time of 50 ms, RF lens (%) of 30, and a normalized AGC target (%) of 250, equivalent to 1e6 ions. Precursor ions with charge states 2- 5 were isolated in the quadrupole with an isolation window of 0.5 m/z. HCD MS2 scans were acquired in a data-dependent manner using a fixed normalized collision energy of 25% and an AGC target (%) of 250, equivalent to 2.5e4 ions, and collected in the ion trap from 150-1,350 m/z, with a maximum injection time of 14 ms and a dynamic exclusion period set to 10 seconds.

*Data Analysis.* For shotgun proteomics data, raw proteomic data files were processed by MaxQuant version 2.4.7.0 (*81*). The UniProt database of reviewed proteins and isoforms from Homo sapiens was retrieved on December 13, 2023. Default MaxQuant parameters were used for processing, along with the following parameters: label-free quantification (LFQ) calculated with a minimum ratio of 1; match between runs enabled; MS/MS not required for LFQ comparisons. In the generated MaxQuant data output, protein identifications were removed that were indicated to be identified by site only, corresponded to reverse sequences, and/or to be potential contaminants by MaxQuant. Protein identifications that generated an intensity value of zero in 50% or more of the analyzed samples were also removed. Missing quantitative values among the remaining protein groups were imputed, log2- transformed, and statistically analyzed using Argonaut (*82*).

For N-glycoproteomics data, raw proteomic data files were processed by MSFragger-Glyco (v20.0) (*83*), using the UniProt database of reviewed proteins and isoforms from Homo sapiens retrieved August 31, 2023. Default search parameters for “Glyco-N-LFQ” workflow were used, except the range for peptides was expanded to 65 amino acids and max mass of 6,500; MaxLFQ quantification and match-between-runs were enabled. The number of missing values (zeroes) were counted across the samples in each condition; glycan-bearing peptide IDs were kept if there were intensity values above zero in at least three values within a single condition. Missing quantitative values among the remaining protein groups were imputed, log2-transformed, and statistically analyzed using Argonaut5.

### Quantitative RT-PCR analysis

Approximately 48 hours before treatment, fibroblasts were seeded in 24-well plates at a density of 1.5 × 10^4^ per well. Cells were treated for 12 hours with WNT3A or control conditioned media as indicated.

Cells were harvested in 800 µl of TRIzol Reagent (Thermo Fisher Scientific Cat. # 15596018) and RNA prepared following manufacturer protocol. 250 ng of RNA was used to synthesize cDNA using iScript Reverse Transcription Supermix (Bio-Rad Laboratories Cat. # 170–8841) following manufacturer protocol. cDNA was diluted 1:100 in water, and 5 µl were mixed with 5 µl of *Power* SYBR™ Green PCR Master Mix (Applied Biosystems Cat. # 4367659) containing 200 nM each of forward and reverse primer for *AXIN2* (Fwd: 5’-GTCCAGCAAAACTCTGAGGG-3’, Rev: 5’-CTGGTGCAAAGACATAGCCA-3’), *TNFRSF19* (Fwd: 5’- GGTGCATTCTGCAGCCAGTCTT-3’, Rev: 5’- CAGGCATCTGAAAACTCGCCAC- 3’), *GAPDH* (Fwd: 5’- AAAGGGTCATCATCTCTG-3’, Rev: 5’- GCTGTTGTCATACTTCTC-3’). Triplicate reactions for each cDNA and primer pair were run on a QuantStudio 5 Real-Time PCR System (Thermo Fisher Scientific) and transcript levels relative to *GAPDH* were calculated using the ΔCt method.

### Preparation of rough microsomes

The original method (*84*) for isolation of microsomal membranes for cotranslational protein translocation from canine pancreas has been adapted to HEK293T cells (*60*). HEK293T cells were grown to ∼80% confluency in 15 cm tissue culture-treated dishes, washed thrice with ∼20 mL ice-cold PBS and collected by scraping in 5-10 mL of PBS. Cells were centrifuged for 5 min at 500 x g and resuspended in 2 mL lysis buffer (10 mM HEPES-NaOH pH 7.4, 250 mM sucrose, 2 mM MgCl_2_, 0.5 mM DTT, 1x SIGMAFAST protease inhibitor cocktail) per dish of cells. Cells were homogenized using a chilled and equilibrated Isobiotec Cell Homogenizer (5–10 single passes, 14 μm clearance) on ice and lysate was cleared twice (1,500 x g for 3 min at 4°C). Microsomes were pelleted (10,000 x g for 10 min at 4°C) and resuspended in microsome buffer (10 mM HEPES-NaOH pH 7.4, 250 mM sucrose, 1 mM MgCl_2_, 0.5 mM DTT) at a density of 1 mL buffer for every 3-4 dishes of cells. Each 1 mL aliquot of microsomes was treated for 10 min at 37°C with 4000 U micrococcal nuclease (New England Biolabs), 2 U RNase-free DNase (Promega), 1 mM CaCl_2_, and 0.5 mM PMSF (Sigma-Aldrich), followed by quenching with 2 mM EGTA. Microsomes were pelleted (10,000 x g for 10 min at 4°C), resuspended in 1 mL microsome buffer containing 40 U SUPERaseIn and 0.1 mM EGTA, and pelleted again (10,000 x g for 10 min at 4°C). The membrane pellet was resuspended in fresh microsome buffer and adjusted to an absorbance at 260 nm of 50-75 and used fresh in translation reactions.

### *In vitro* translation (IVT) and glycosylation analysis

RNA encoding indicated HSP90B1 nascent chains were prepared from PCR amplified and purified DNA template containing a T7 promoter and carried out at 37°C for 2 hours using the HiScribe® T7 High Yield RNA Synthesis Kit (New England Biolabs) and purified using the RNeasy Mini Kit (Qiagen). For *in vitro* glycosylation analysis of full length protein products, 50 μL IVT reactions were carried out for 2 hours at 30°C using commercially available Nuclease-Treated Rabbit Reticulocyte Lysate (RRL, Promega) and reaction components were added in the following order: 35 μL RRL, 1 μL of 1 mM complete amino acid mixture (combine amino acid mixtures provided with RRL), 5 μL microsomal membranes, 2 μg RNA, and nuclease-free water. IVT reaction was added immediately to 200 μL of immunoprecipitation buffer (50 mM Tris pH 7.4, 150 mM NaCl, 1% NP-40, 10% glycerol) containing 10 μL anti-FLAG M2 resin. For *in vitro* glycosylation analysis of translation intermediates, 200 μL IVT reactions were set up as described above and 50 μL was removed at each time point and added to 200 μL immunoprecipitation buffer containing 0.5 mM puromycin and 10 μL anti-FLAG M2 resin. Reactions were incubated with anti-FLAG M2 resin for 2 hours at 4°C, washed 3 times with immunoprecipitation buffer, and eluted twice in immunoprecipitation buffer containing 1 mg/mL 3xFLAG peptide (Sigma-Aldrich) (25 μL at 4°C overnight and 25 μL for 30 min at room temperature). Eluates were combined, applied to equilibrated Ultrafree-MC Centrifugal Filter (MilliporeSigma), and spun at 12,000 x g for 4 min at 4°C. Half of eluate was treated with Endo H prior to immunoblot analysis.

### *In vitro* translation (IVT) and immunoprecipitation of ribosome nascent chain complexes

Templates for synthesis of stalled HSP90B1 nascent chains were PCR amplified using reverse primers encoding a terminal Met-Leu-Lys-Val (5′-CACCTTGAGCAT-3’) sequence and lacking a stop codon. RNA was prepared as described above. Translation reactions of 500 μL containing 60 μL microsomal membranes was prepared as described above and carried out for 1 hour at 30°C. Reactions were immediately diluted in cold 500 μL IVT stop buffer (50 mM HEPES pH 7.4, 200 mM NaCl, 10 mM MgCl_2_) and centrifuged at 12,500 x g for 10 min, 4°C. Membrane pellet was washed once with 1 mL IVT stop buffer and centrifuged at 12,500 x g for 10 min, 4°C. Membrane pellet was resuspended in 500 μL IVT stop buffer and treated for 20 min at room temperature with 5000 U micrococcal nuclease, 1 mM CaCl_2_, and 0.6 mM PMSF, followed by quenching with 2.5 mM EGTA, and pelleted at 12,500 x g for 10 min, 4°C. Membrane pellet was solubilized in 200 μL solubilization buffer (50 mM HEPES-KOH pH 7.4, 250 mM sucrose, 250 mM KOAc, 10 mM MgCl_2_, 2.5% digitonin) for 45 min, rotating at 4°C, followed by addition of 200 μL of dilution buffer (50 mM HEPES-KOH pH 7.4, 250 mM sucrose, 150 mM KOAc, 10 mM MgCl_2_), and cleared at 12,500 x g for 15 min, 4°C. Cleared supernatant was incubated overnight with 15 μL anti-FLAG M2 resin. Resin was washed thrice with 400 μL wash buffer (50 mM HEPES-KOH pH 7.4, 250 mM sucrose, 200 mM KOAc, 10 mM MgCl_2_, 0.4% digitonin) and eluted twice in IVT stop buffer containing 1 mg/mL 3xFLAG peptide and 0.4% digitonin by rotating for 30 min at 4°C. Eluates were combined, applied to equilibrated Ultrafree-MC Centrifugal Filter (MilliporeSigma), and spun at 12,000 x g for 4 min at 4°C. For RNaseA and Endo H treatments, 15 μL of eluate was first treated with 50 μg/mL RNaseA, 10 mM EDTA, 0.5% SDS in a total of 20 μL for 15 min at 37°C. Then 1 μL of 1 M DTT was added and the sample boiled for 10 min, then 1 μL Endo H and 2.5 μL GlycoBuffer 3 (New England Biolabs) was added and incubated for 1 hour at 37°C, followed by immunoblot analysis.

### Pulse chase analysis

Clonally derived *HSP90B1^-/-^* and *HSP90B1^-/-^*; *CCDC134^-/-^* RKO cells stably expressing 3xFLAG- HSP90B1 were seeded at a density of 2 × 10^6^ per 6 cm tissue culture-treated dish and grown for two days. Cells were washed twice with PBS and changed into starvation medium: Met/Cys-free DMEM (Gibco Cat. # 21013024) containing 10% dialyzed FBS for 40 min. For pulse labeling, cells were changed into 1 mL of starvation media containing 200 μCi/mL EasyTag EXPRESS^35^S Protein Labeling Mix (PerkinElmer NEG772002MC), grown for indicated time points, immediately changed into ice-cold PBS and collected on ice. For pulse chase analysis, cells were pulse labeled as described and immediately changed into chase medium (complete DMEM containing an additional 2 mM Met and 2 mM Cys and 100 μg/mL cycloheximide) for the indicated time points and collected in ice-cold PBS. Whole cell lysate was prepared in lysis buffer (50 mM Tris at pH 7.4, 150 mM NaCl, 1% NP-40, 0.25%

Deoxycholate, 10% glycerol, 1x SIGMAFAST protease inhibitor cocktail (MilliporeSigma), and 1x PhosSTOP phosphatase inhibitor cocktail (Roche)) and a small aliquot was used in LSC analysis to ensure equivalent CPMs of lysates were used for immunoprecipitation (generally 300-800 μg). Lysates were incubated with 10 μL anti-FLAG M2 resin for 2 hours at 4°C and washed 3 times with lysis buffer. 25% of the sample was used for Endo H (New England Biolabs) treatment following manufacturer protocol. All samples were eluted in NuPAGE-LDS sample buffer containing 100 mM DTT or 50 mM TCEP for SDS-PAGE analysis and transferred onto PVDF membrane (Bio-Rad Laboratories Cat. #162- 0177). Dried membranes were exposed to a phosphor screen (Molecular Dynamics), scanned on Typhoon FLA 9500, and quantified in ImageStudio (LI-COR) to determine the distribution of glycoforms.

### Evolutionary sequence analysis

Human CCDC134 9 (NP_079097.1) was extracted from the National Center for Biotechnology Information (NCBI) Genbank database. Sequence similarity searches were performed using it as the query with the PSI-BLAST program against the NCBI non-redundant (nr) database or the same database clustered down to 50% sequence identity using the MMseqs program with a profile-inclusion threshold was set at an e-value of 0.01. Profile-profile searches were performed with the HHpred program. Multiple sequence alignments (MSAs) were constructed using the MAFFT program. Sequence logos were constructed using these alignments with the ggseqlogo library for the R language. Signal peptides were predicted using a deep neural network as implemented in SignalP 6.

### Evolutionary structure analysis

The JPred program was used to predict secondary structures using MSAs (see above). Sequence complexity analysis was performed using the SEG program. Structural models were generated using the RoseTTAfold and Alphafold2 programs. Multiple alignments of related sequences (>30% similarity) were used to initiate HHpred searches to be used by the neural networks deployed by these programs.

Structures were rendered, compared, and superimposed using the Mol* program.

### Comparative genomics and phylogenetic analysis

Clustering of protein sequences was performed using MMSEQS with empirical adjustment of the length of aligned regions and bit-score density threshold. Phylogenetic analysis was performed using the maximum-likelihood method with the WAG substitution matrix and 20 gamma-distributed categories with the IQTree program.

### Statistical analysis

Data analysis and visualization were performed in GraphPad Prism 10. Model figure 6B was made with Biorender.com and all other figures were made in Adobe Illustrator 2022. The predicted human CCDC134 structure was generated using AlphaFold.

The one-way ANOVA test with Sidak’s multiple comparison or Dunnett’s multiple comparison were used to compare three or more groups with one independent variable. A two-way ANOVA test with Dunnett’s multiple comparison or Tukey’s multiple comparison were used to compare three or more groups with two independent variables. All comparisons shown were prespecified. All experiments were performed at least three different times with similar results. We note that a small sample size (*n*=3) makes it difficult to assess whether the variance between different samples is comparable. Throughout the paper, the *p*-values for the comparisons from GraphPad Prism 10 are denoted on on the graphs according to the following key: **** *p*-value<0.0001, *** *p*-value<0.001, ** *p*-value<0.01, * *p*-value<0.05, and n.s. Non-significant. Descriptions of replicates are included in figure legends.

Genome-wide loss-of-function CRISPR/Cas9 screens were performed twice under independent conditions and the duplicates from each screen were analyzed together using the MAGeCK tool (**Fig.1B**). Mass spectrometry analysis used six individual replicates (each replicate is an individual cell pellet processed in a different mass spectrometry run) and error bars depict standard error of the mean where indicated.

---

**Supplementary Figure 1.**
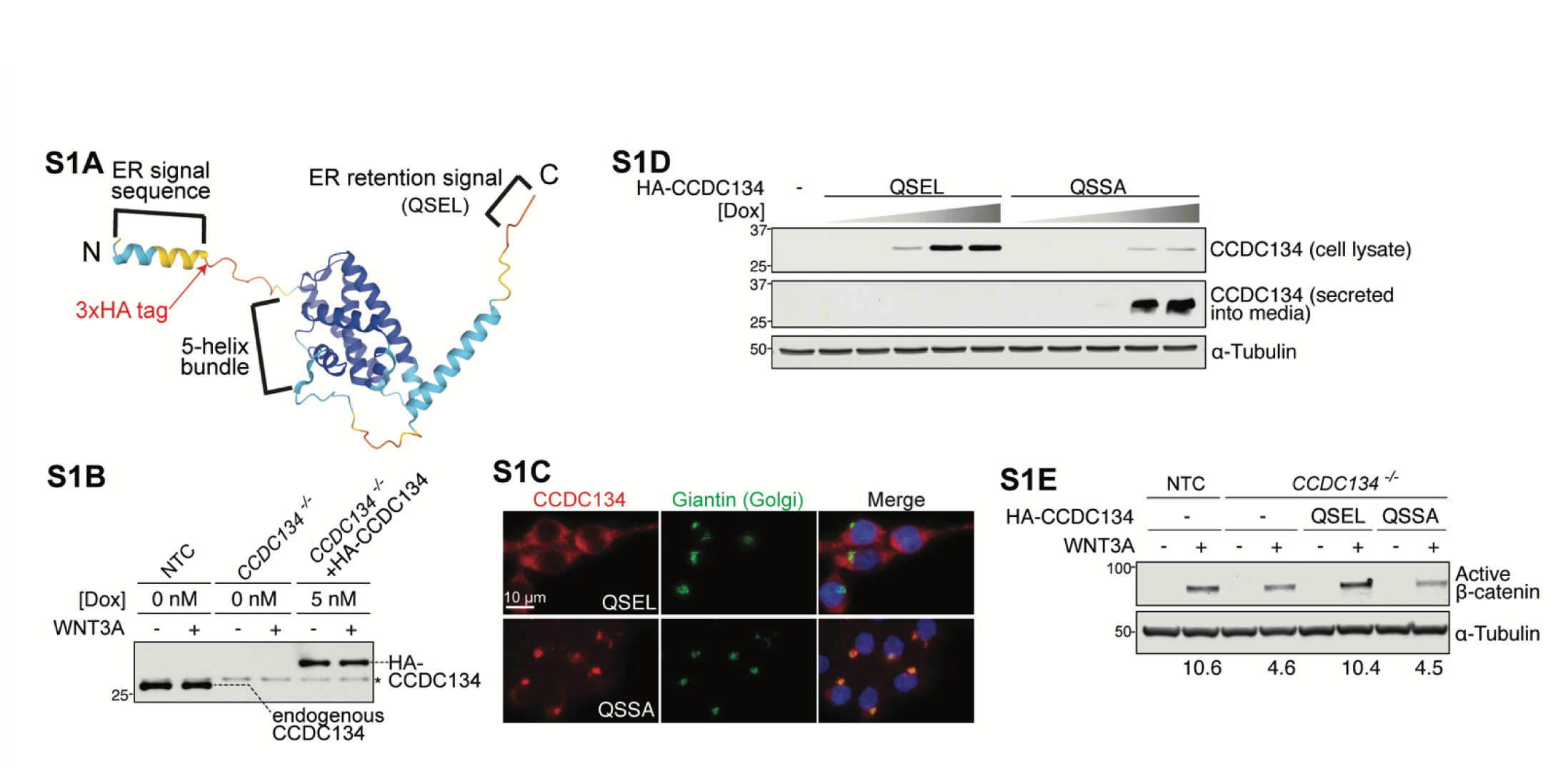
CCDC134 is a protein resident in the ER. **(A)** AlphaFold prediction of the structure adopted by CCDC134. ER signal sequence (characterized in (*23*)) and ER retrieval signal (QSEL; characterized here) are labeled. The position of the 3xHA sequence used to epitope tag CCDC134 in most of our experiments is noted with a red arrowhead. **(B)** Abundance of 3xHA-CCDC134 in cells stably expressing a doxycycline (Dox)-inducible transgene after treatment with 5 nM Dox was measured by immunoblotting with an anti-CCDC134 antibody. Abundance of endogenous CCDC134 in control (NTC) cells is shown as a comparison. These conditions were used throughout our experiments to achieve near-endogenous levels of 3xHA-CCDC134. **\*** marks a nonspecific band. **(C)** Fluorescence microscopy was used to assess the subcellular localization of 3xHA-CCDC134 (red) carrying its native QSEL ER retention sequence or a mutant QSSA sequence at the C-terminus after stable expression in *CCDC134^-/-^* cells. Giantin (green) was used to mark the Golgi. Wild-type CCDC134 was found in a reticular pattern throughout the cytoplasm, consistent with ER localization (top row), but the CCDC134-QSSA mutant was mostly localized in the Golgi, a hallmark of secreted proteins (bottom row)(*26*). Scale bar: 10 μm. **(D)** Abundance of 3xHA-CCDC134 carrying the native QSEL or mutant QSSA ER retention sequence found in cells or secreted into media. The CCDC134 variants were stably expressed in *CCDC134^-/-^* cells under the control of a doxycycline-inducible promoter. Cells and accompanying media were harvested for immunoblotting 24 hours after treatment with increasing concentrations of doxycycline (0, 2, 10, 50, 250 nM). **(E)** Active β-catenin abundance (+/-WNT3A) in control or *CCDC134^-/-^* cells stably expressing the QSEL or QSSA variants of 3xHA-CCDC134 (see **C**) under a doxycycline-inducible promoter. CCDC134 variants were induced using a doxycycline concentration (5 nM) that resulted in near-endogenous protein abundance (see **S1B)**. Numbers below the lanes show the WNT3A-induced fold-change in active β-catenin abundance normalized to α-Tubulin abundance.

**Supplementary Figure 2.**
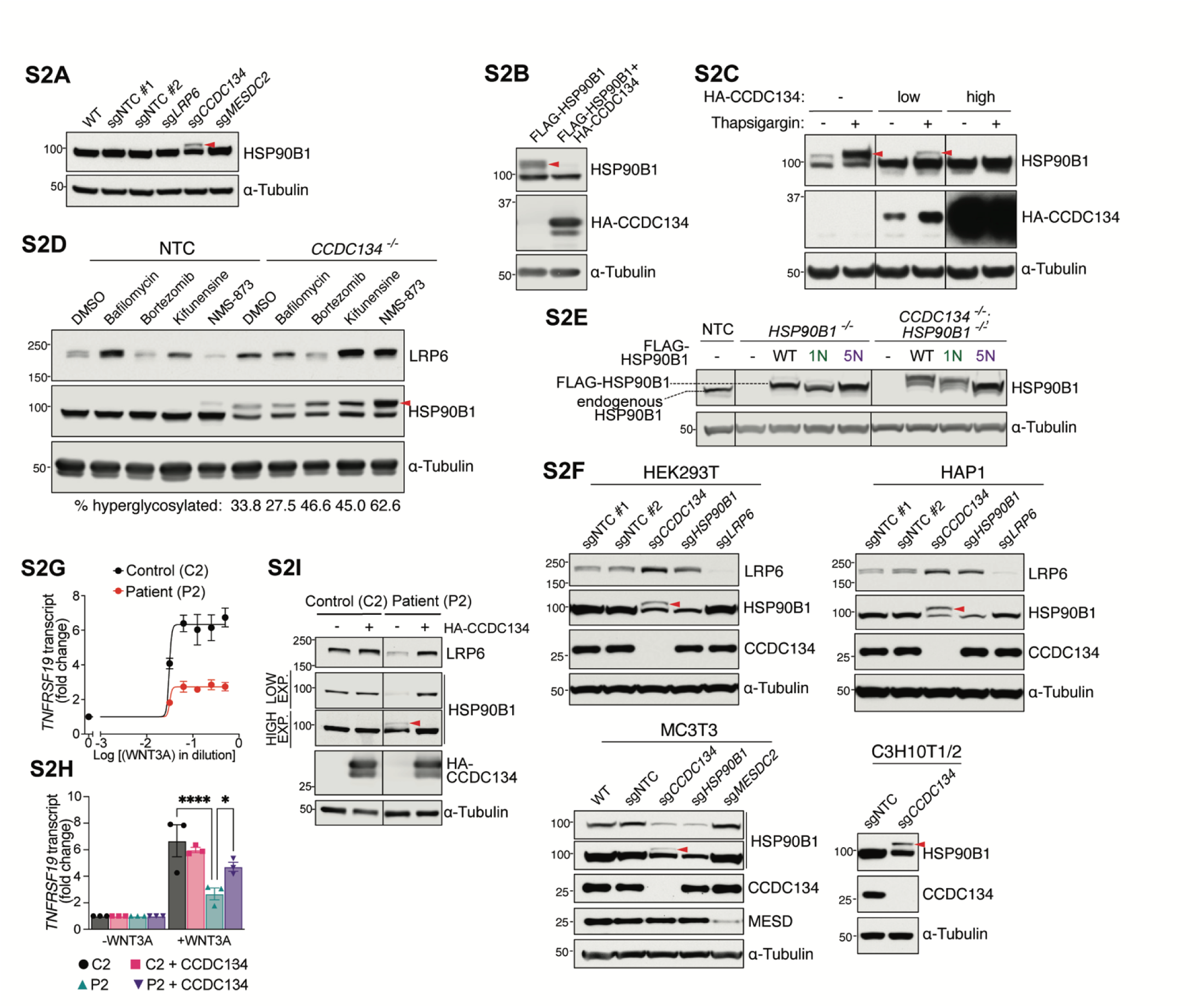
CCDC134 suppresses the hyperglycosylation and subsequent degradation of HSP90B1 via the endoplasmic-reticulum-associated protein degradation (ERAD) pathway. (A) Abundance and glycosylation status of HSP90B1 on SDS-PAGE in lysates from WT RKO cells, cells expressing a control (NTC) sgRNA or sgRNAs targeting the indicated genes. The hyperglycosylated HSP90B1 band is indicated with a red arrowhead. (B) Impact of CCDC134 co-expression on HSP90B1 hyperglycosylation (red arrowhead) when the indicated proteins are transiently expressed in HEK293T cells. (C) Effects of low and high 3xHA-CCDC134 expression (from a stably integrated, doxycycline-inducible transgene) on HSP90B1 hyperglycosylation (red arrowhead) in *CCDC134^-/-^*RKO cells treated with 0.1 uM thapsigargin for 24 hrs. Low (near-endogenous, see **fig.S1B**) and high CCDC134 expression was achieved by supplementing media with 5 nM or 25 nM doxycycline, respectively. (D) Abundances and glycosylation status of HSP90B1 and LRP6 in lysates from clonally derived control (NTC) and *CCDC134^-/-^* RKO cells treated (18 hours) with small molecule inhibitors of (1) lysosomal acidification (50 nM Bafilomycin A1), (2) the proteasome (1 μM bortezomib), or (3) the ERAD pathway (5 μg/mL kifunensine or 5 μM NMS-873). As a control, media was also supplemented with an equivalent concentration of the solvent (DMSO) contributed by the inhibitor stock solutions. The hyperglycosylated HSP90B1 band is indicated with a red arrowhead. Numbers below the lanes show the percentage of HSP90B1 that is hyperglycosylated; note the increase in bortezomib, kifunensine, and NMS-873 treatments. (E) Abundances of endogenous HSP90B1 in control (NTC) cells compared to stably integrated 3xFLAG- HSP90B1 variants in *HSP90B1^-/-^* and *CCDC134^-/-^*;*HSP90B1^-/-^*cell lines (see **Fig.2H, 2I**). (F) LRP6 abundance and HSP90B1 hyperglycosylation (red arrowhead) in a panel of human (HEK293T, HAP1) and mouse (MC3T3, C3H10T1/2) cell lines expressing sgRNAs targeting the indicated genes (NTC=non-targeting control sgRNA). (G) Expression of the WNT target gene *TNFRSF19* (mean +/- SEM of *TNFRSF19* mRNA normalized to *GAPDH* mRNA measured by qRT-PCR in three independent experiments) was quantified in primary patient fibroblasts treated with 0, 3.125, 6.25, 12.5, 25, 50% WNT3A conditioned media. See **Fig.2L** for the same comparison using a different WNT target gene (*AXIN2*). (H) Expression of the WNT target gene *TNFRSF19* (mean +/- SEM of *TNFRSF19* mRNA normalized to *GAPDH* mRNA, measured by qRT-PCR in three independent experiments) in primary patient fibroblasts stably expressing 3xHA-CCDC134 treated with 25% WNT3A conditioned media. Statistical significance was determined by two-way ANOVA Dunnett’s multiple comparisons test; * p<0.05, **** p<0.0001. See **Fig.2M** for the same comparison using a different WNT target gene (*AXIN2*). (I) Abundances of LRP6 and HSP90B1 in primary fibroblasts isolated from a *CCDC134^-/-^* patient (P2, **Fig.2J**) with or without stable re-expression of 3xHA-CCDC134 (*5*).

**Supplementary Figure 3.**
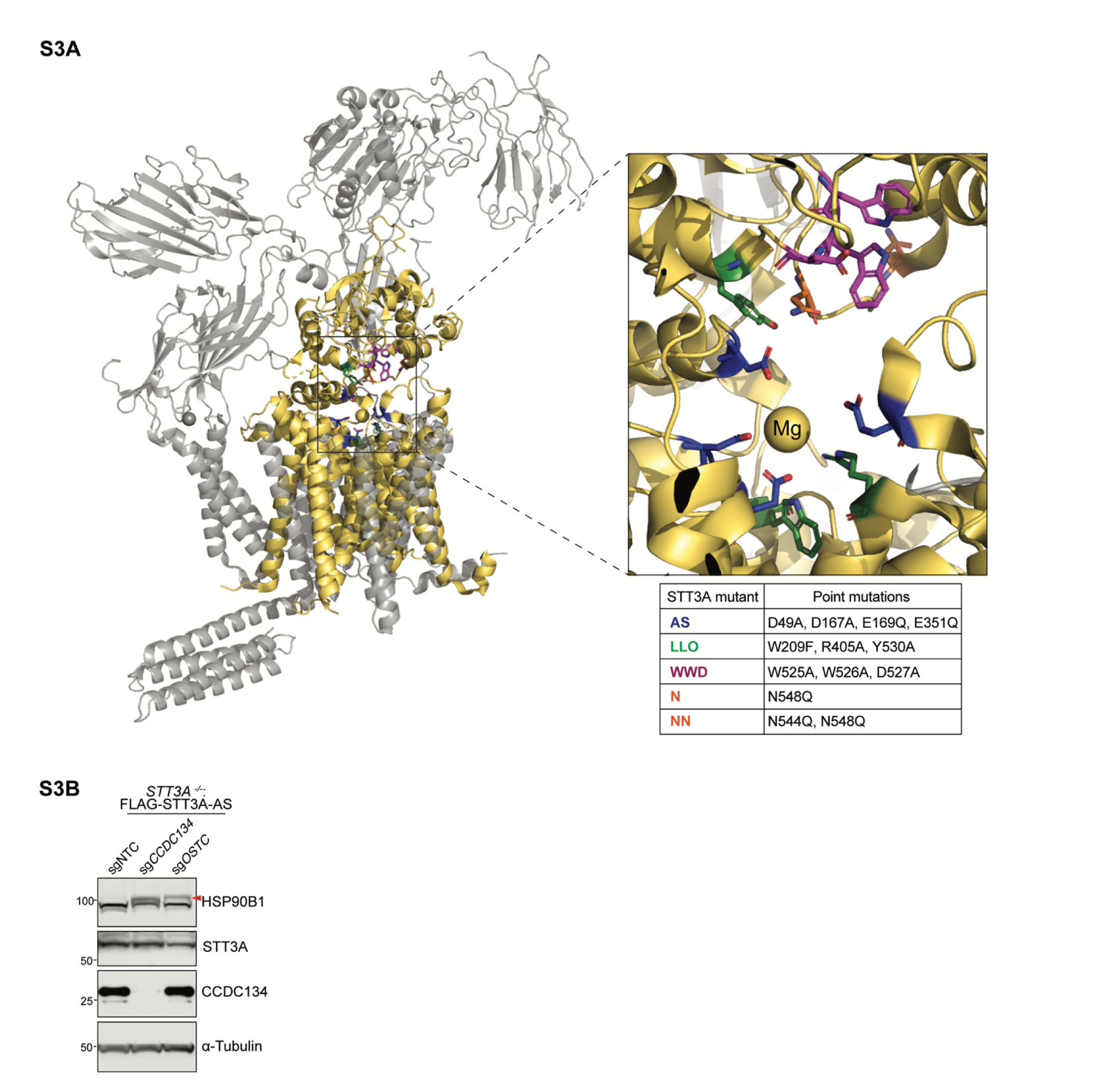
Catalytic activity of STT3A is not required for its role in regulating HSP90B1 hyperglycosylation. (A) Cryo-EM structure of the OST-A complex (PDB 6S7O (*11*)) with STT3A highlighted in yellow. Residues that were mutated to generate the STT3A variants tested in **Fig.3H** and **3I** are shown on the structure and listed in the table below. Variants carry mutations in residues involved in active site (AS, blue) chemistry, lipid-linked oligosaccharide binding (LLO, green), sequon binding (WWD, magenta) or N-glycosylation of STT3A itself (N and NN, orange). (B) Abundance and glycosylation status (red arrowhead) of HSP90B1 in *STT3A^-/-^* cells stably expressing (1) catalytically inactive FLAG-STT3A carrying mutations in its active site (AS, see **A**) and (2) sgRNAs targeting CCDC134 or OSTC. NTC=non-targeting control sgRNA. See also **Fig.3**.

**Supplementary Figure 4.**
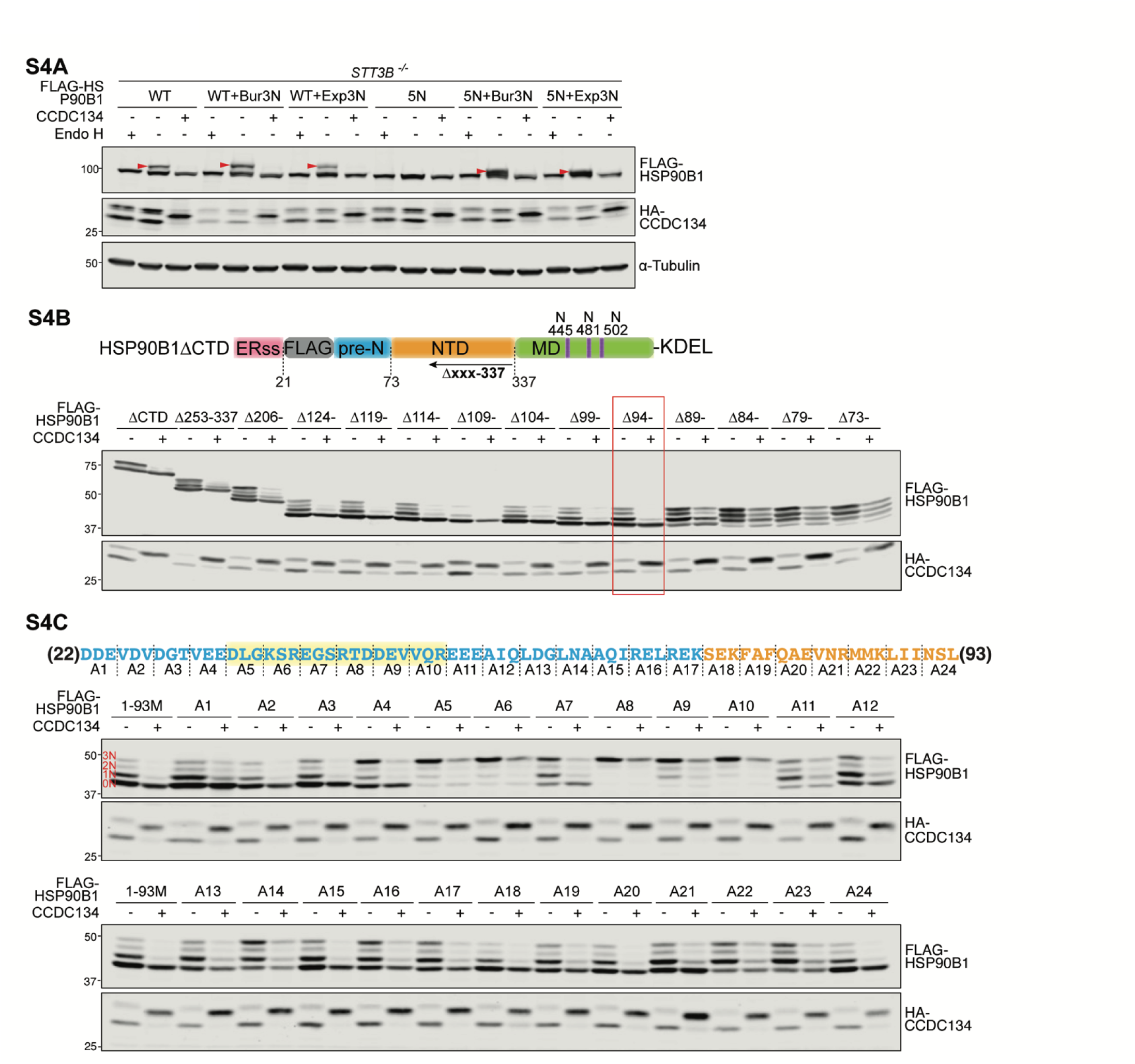
Identification of the region of HSP90B1 that regulates its own N-glycosylation using truncation analysis and alanine scanning mutagenesis. (A) Glycosylation status of full-length FLAG-HSP90B1 variants co-expressed in *STT3B^-/-^* HEK293T cells with functional CCDC134 (**+**) carrying a native ER signal sequence or non-functional CCDC134 (-) lacking an ER signal sequence. WT refers to HSP90B1 carrying all native sequons while 5N refers to a variant that lacks the five facultative sequons (shown in **Fig.2E**). Bur3N and Exp3N refer to mutations that introduce three additional artificial sequons predicted to be buried (Bur) or exposed (Exp) based on the PDB 5ULS HSP90B1 structure. Red arrowheads indicate hyperglycosylated forms of each variant. (B) Glycosylation status of the indicated deletion mutants of FLAG-HSP90B1 lacking its CTD and containing only three sequons in the M domain (ΔCTD, see **Fig.4A**). The deletion series starts at amino acid 337 and extends sequentially into the NTD and pre-N segments. The minimum construct that still retains regulation of glycosylation is boxed in red, this is also named the 1-93M construct. (C) Triplet alanine scanning mutagenesis was used to identify amino acid residues within the pre-N domain of HSP90B1 that regulates its own glycosylation. Sets of three consecutive residues (A1-A24, as shown in the pre-N sequence above the immunoblots) were mutated to Ala-Ala-Ala in the 1-93M variant of HSP90B1 (see **Fig.4A** for domain structure) and glycosylation tested by transiently transfecting the encoding constructs into HEK293T cells. See also **Fig.4**.

**Supplementary Figure 5.**
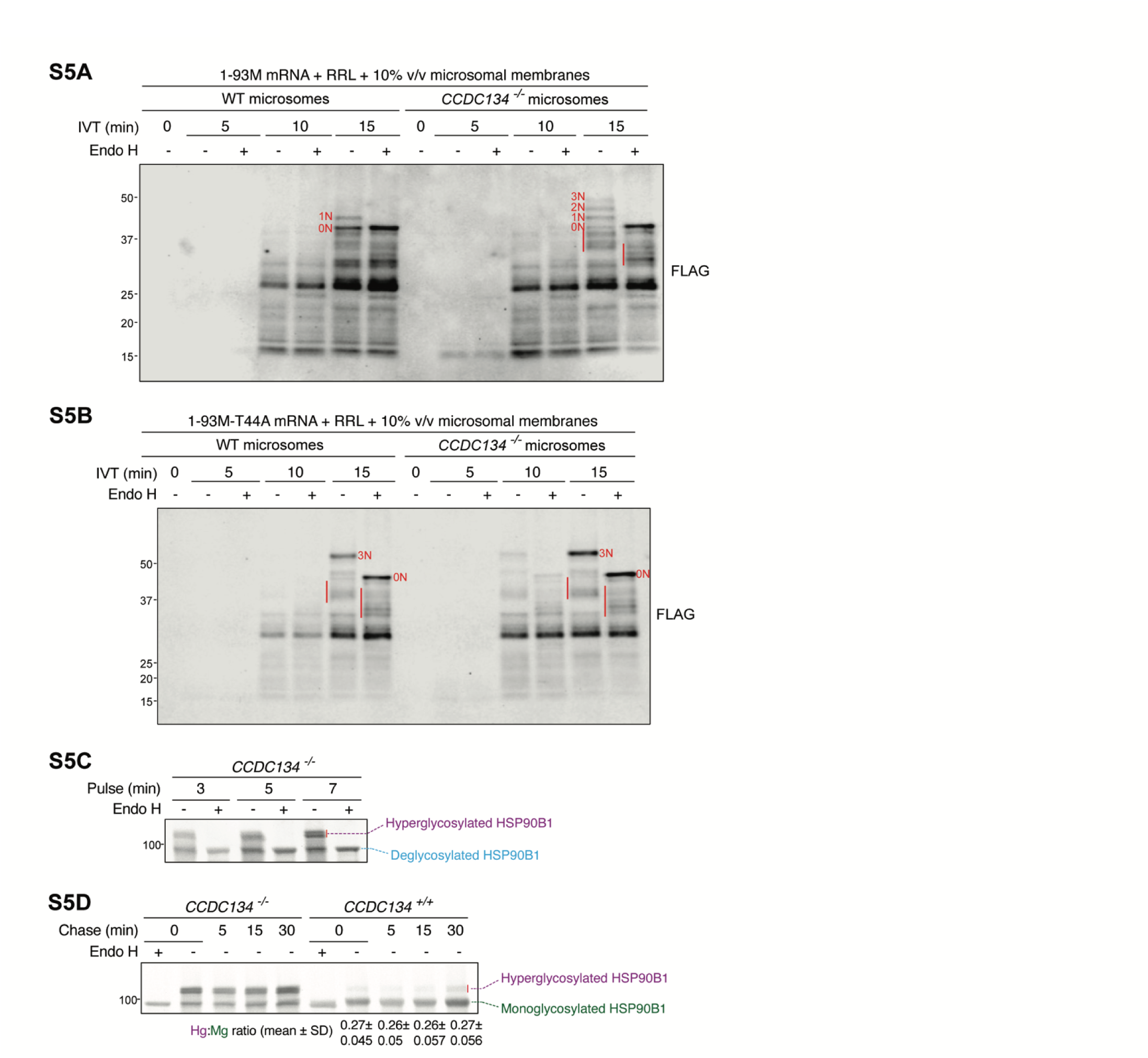
Co-translation hyperglycosylation of HSP90B1. (**A, B**) Translation time course of 1-93M (**A**) or 1-93M carrying the T44A mutation in the “SRT” pseudosubstrate site (**B**) in rabbit reticulocyte lysate (RRL) supplemented with rough microsomes isolated from wild-type or *CCDC134^-/-^* HEK293T cells. The sizes and Endo H sensitivities of the translation products were analyzed after anti-FLAG immunoprecipitation. Glycosylated translational intermediates (highlighted by a red line) were identified by their size (shorter than the full-length protein seen at 15 min) and sensitivity to Endo H. (C) Pulse-labeling of HSP90B1 in clonal *HSP90B1^-/-^*;*CCDC134^-/-^* double knock-out cells stably expressing 3xFLAG-HSP90B1. Culture media was supplemented with ^35^S-Methionine and ^35^S-Cysteine for 3, 5 or 7 minutes, followed by lysis and immunoprecipitation of HSP90B1 on anti-FLAG beads. The Endo H-treated sample is equivalent to 33% of the undigested sample. (D) Pulse labeling (5 min) of HSP90B1 with ^35^S-Methionine/^35^S-Cysteine in *HSP90B1^-/-^* or *HSP90B1^-/-^* ;*CCDC134^-/-^* cells stably expressing 3xFLAG-HSP90B1 was followed by a chase in unlabelled media for 5, 10 or 30 minutes. The ratio of the hyperglycosylated (Hg) to monoglycosylated (Mg) HSP90B1 band intensity is indicated from three independent experiments.

**Supplementary Figure 6.**
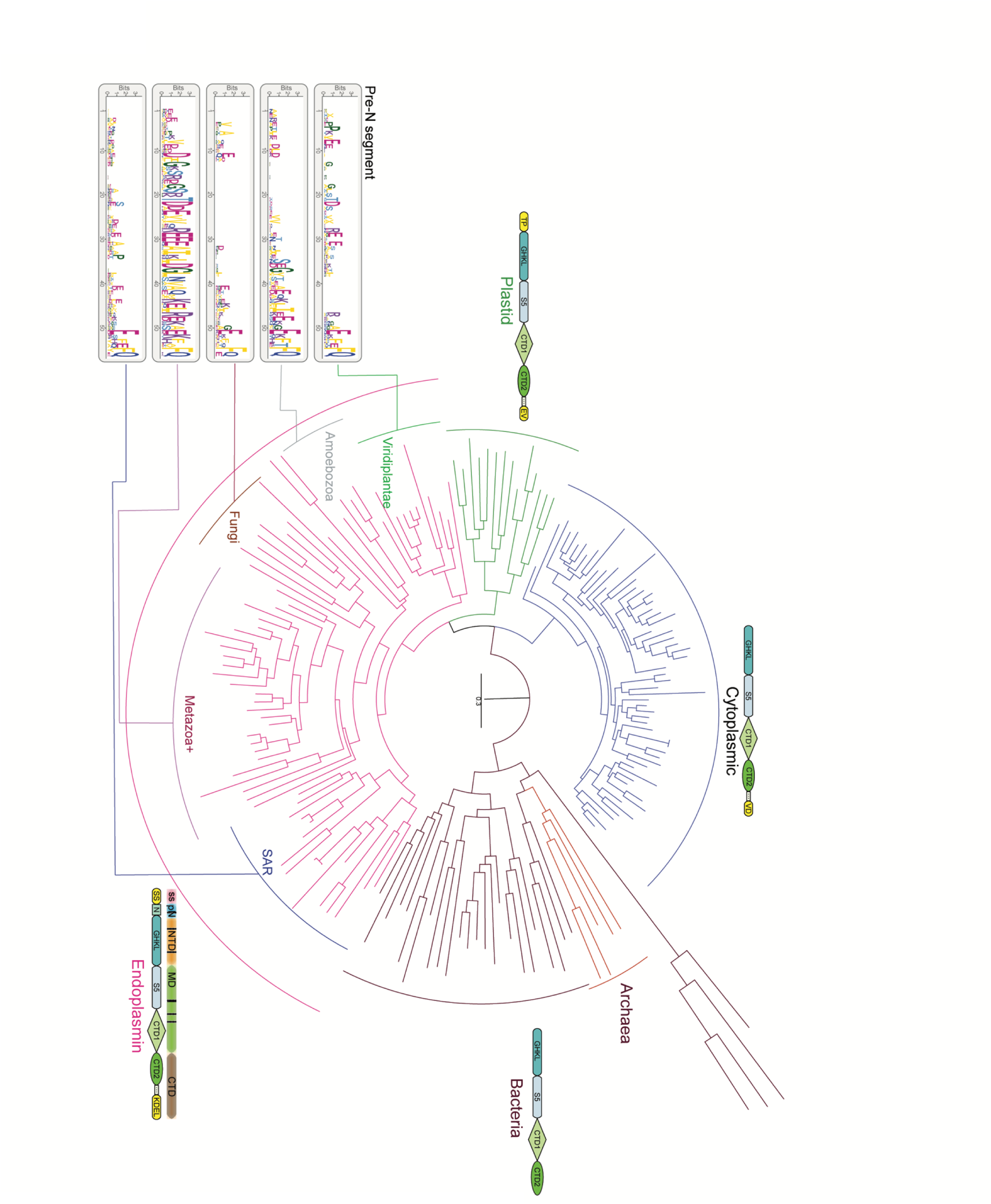
Phylogenetic tree of HSP90. The 4 major clades (prokaryotic, eukaryotic cytoplasmic, eukaryotic plastid, and eukaryotic endoplasmin) are colored distinctly. For each clade the formal domain architecture is indicated. For the endoplasmins, the HSP90B1 domain diagram used throughout this manuscript is juxtaposed for comparison and glycosylation sites and marked as black lines. Within the endoplasmin clade (which includes HSP90B1), the subclades corresponding to major eukaryotic lineages are marked separately. The conservation pattern in the pre-N segment unique to the endoplasmins is shown for each major subclade as sequence logos. The logos are anchored to the start of the HSP90 NTD (or ATP-binding GHKL domain) and the residues are scaled according to the conservation bit score. Metazoa+ indicates the Metazoa+ sister lineages choanoflagellates and Filasterians. Other annotated features include the S5 region, supplies the lysine that stabilizes the hypercharged state during hydrolysis; CTD1 and CTD2, two domains which together are called C-terminal; GHKL, ancient ATP binding domain found in Gyrase, Hsp90, histidine Kinases and MutL; TP, transit peptide in plastid HSP90; EV/VD/KDEL, terminal motifs.

## Notes

### Competing Interest Statement

The authors have declared no competing interest.

## REFERENCES AND NOTES

1. P. Gagneux, T. Hennet, A. Varki, “Biological Functions of Glycans” in Essentials of Glycobiology, A. Varki, R. D. Cummings, J. D. Esko, P. Stanley, G. W. Hart, M. Aebi, D. Mohnen, T. Kinoshita, N. H. Packer, J. H. Prestegard, R. L. Schnaar, P. H. Seeberger, Eds. (Cold Spring Harbor Laboratory Press, Cold Spring Harbor (NY), 2022).

2. C. Ruiz-Canada, D. J. Kelleher, R. Gilmore, Cotranslational and posttranslational N-glycosylation of polypeptides by distinct mammalian OST isoforms. Cell 136, 272–283 (2009).

3. E. A. Ansa-Addo, J. Thaxton, F. Hong, B. X. Wu, Y. Zhang, C. W. Fugle, A. Metelli, B. Riesenberg, K. Williams, D. T. Gewirth, G. Chiosis, B. Liu, Z. Li, Clients and Oncogenic Roles of Molecular Chaperone gp96/grp94. Curr. Top. Med. Chem. 16, 2765–2778 (2016).

4. D. Dersh, S. M. Jones, D. Eletto, J. C. Christianson, Y. Argon, OS-9 facilitates turnover of nonnative GRP94 marked by hyperglycosylation. Mol. Biol. Cell 25, 2220–2234 (2014).

5. J. Dubail, P. Brunelle, G. Baujat, C. Huber, M. Doyard, C. Michot, P. Chavassieux, A. Khairouni, V. Topouchian, S. Monnot, E. Koumakis, V. Cormier-Daire, Homozygous Loss-of-Function Mutations in CCDC134 Are Responsible for a Severe Form of Osteogenesis Imperfecta. J. Bone Miner. Res. 35, 1470–1480 (2020).

6. G. A. Khoury, R. C. Baliban, C. A. Floudas, Proteome-wide post-translational modification statistics: frequency analysis and curation of the swiss-prot database. Sci. Rep. 1 (2011).

7. D. F. Zielinska, F. Gnad, J. R. Wiśniewski, M. Mann, Precision mapping of an in vivo N- glycoproteome reveals rigid topological and sequence constraints. Cell 141, 897–907 (2010).

8. I. J. Chang, M. He, C. T. Lam, Congenital disorders of glycosylation. Ann Transl Med 6, 477 (2018).

9. C. Reily, T. J. Stewart, M. B. Renfrow, J. Novak, Glycosylation in health and disease. Nat. Rev. Nephrol. 15, 346–366 (2019).

10. S. Shrimal, N. A. Cherepanova, R. Gilmore, DC2 and KCP2 mediate the interaction between the oligosaccharyltransferase and the ER translocon. J. Cell Biol. 216, 3625–3638 (2017).

11. A. S. Ramírez, J. Kowal, K. P. Locher, Cryo-electron microscopy structures of human oligosaccharyltransferase complexes OST-A and OST-B. Science 366, 1372–1375 (2019).

12. K. Braunger, S. Pfeffer, S. Shrimal, R. Gilmore, O. Berninghausen, E. C. Mandon, T. Becker, F. Förster, R. Beckmann, Structural basis for coupling protein transport and N-glycosylation at the mammalian endoplasmic reticulum. Science 360, 215–219 (2018).

13. R. Wild, J. Kowal, J. Eyring, E. M. Ngwa, M. Aebi, K. P. Locher, Structure of the yeast oligosaccharyltransferase complex gives insight into eukaryotic N-glycosylation. Science 359, 545– 550 (2018).

14. L. Bai, T. Wang, G. Zhao, A. Kovach, H. Li, The atomic structure of a eukaryotic oligosaccharyltransferase complex. Nature 555, 328–333 (2018).

15. K. J. Colley, A. Varki, R. S. Haltiwanger, T. Kinoshita, Cellular Organization of Glycosylation (Cold Spring Harbor Laboratory Press, 2022).

16. K. T. Schjoldager, Y. Narimatsu, H. J. Joshi, H. Clausen, Global view of human protein glycosylation pathways and functions. Nat. Rev. Mol. Cell Biol. 21, 729–749 (2020).

17. N. M. Riley, A. S. Hebert, M. S. Westphall, J. J. Coon, Capturing site-specific heterogeneity with large-scale N-glycoproteome analysis. Nat. Commun. 10, 1311 (2019).

18. L. F. Zacchi, B. L. Schulz, N-glycoprotein macroheterogeneity: biological implications and proteomic characterization. Glycoconj. J. 33, 359–376 (2016).

19. R. Nusse, H. Clevers, Wnt/β-Catenin Signaling, Disease, and Emerging Therapeutic Modalities. Cell 169, 985–999 (2017).

20. A. M. Lebensohn, R. Dubey, L. R. Neitzel, O. Tacchelly-Benites, E. Yang, C. D. Marceau, E. M. Davis, B. B. Patel, Z. Bahrami-Nejad, K. J. Travaglini, Y. Ahmed, E. Lee, J. E. Carette, R. Rohatgi, Comparative genetic screens in human cells reveal new regulatory mechanisms in WNT signaling. Elife 5, e21459 (2016).

21. B. Liu, M. Staron, F. Hong, B. X. Wu, S. Sun, C. Morales, C. E. Crosson, S. Tomlinson, I. Kim, D. Wu, Z. Li, Essential roles of grp94 in gut homeostasis via chaperoning canonical Wnt pathway. Proc. Natl. Acad. Sci. U. S. A. 110, 6877–6882 (2013).

22. J.-C. Hsieh, L. Lee, L. Zhang, S. Wefer, K. Brown, C. DeRossi, M. E. Wines, T. Rosenquist, B. C. Holdener, Mesd encodes an LRP5/6 chaperone essential for specification of mouse embryonic polarity. Cell 112, 355–367 (2003).

23. J. Huang, T. Shi, T. Ma, Y. Zhang, X. Ma, Y. Lu, Q. Song, W. Liu, D. Ma, X. Qiu, CCDC134, a novel secretory protein, inhibits activation of ERK and JNK, but not p38 MAPK. Cell. Mol. Life Sci. 65, 338–349 (2008).

24. B. Yu, T. Zhang, P. Xia, X. Gong, X. Qiu, J. Huang, CCDC134 serves a crucial role in embryonic development. Int. J. Mol. Med. 41, 381–390 (2018).

25. S. Yin, Q. Liao, Y. Wang, Q. Shi, P. Xia, M. Yi, J. Huang, Ccdc134 deficiency impairs cerebellar development and motor coordination. Genes Brain Behav. 20, e12763 (2021).

26. I. Raykhel, H. Alanen, K. Salo, J. Jurvansuu, V. D. Nguyen, M. Latva-Ranta, L. Ruddock, A molecular specificity code for the three mammalian KDEL receptors. J. Cell Biol. 179, 1193–1204 (2007).

27. J. Huang, L. Zhang, W. Liu, Q. Liao, T. Shi, L. Xiao, F. Hu, X. Qiu, CCDC134 interacts with hADA2a and functions as a regulator of hADA2a in acetyltransferase activity, DNA damage-induced apoptosis and cell cycle arrest. Histochem. Cell Biol. 138, 41–55 (2012).

28. H. H. Freeze, C. Kranz, Endoglycosidase and glycoamidase release of N-linked glycans. Curr. Protoc. Protein Sci. **Chapter** 12, Unit12.4 (2010).

29. A. L. Tarentino, T. H. Plummer Jr, Enzymatic deglycosylation of asparagine-linked glycans: purification, properties, and specificity of oligosaccharide-cleaving enzymes from Flavobacterium meningosepticum. Methods Enzymol. 230, 44–57 (1994).

30. N. A. Cherepanova, S. V. Venev, J. D. Leszyk, S. A. Shaffer, R. Gilmore, Quantitative glycoproteomics reveals new classes of STT3A- and STT3B-dependent N-glycosylation sites. J. Cell Biol. 218, 2782–2796 (2019).

31. R. A. Mazzarella, M. Green, ERp99, an abundant, conserved glycoprotein of the endoplasmic reticulum, is homologous to the 90-kDa heat shock protein (hsp90) and the 94-kDa glucose regulated protein (GRP94). J. Biol. Chem. 262, 8875–8883 (1987).

32. D. Qu, R. A. Mazzarella, M. Green, Analysis of the structure and synthesis of GRP94, an abundant stress protein of the endoplasmic reticulum. DNA Cell Biol. 13, 117–124 (1994).

33. S. E. Cala, GRP94 hyperglycosylation and phosphorylation in Sf21 cells. Biochim. Biophys. Acta 1496, 296–310 (2000).

34. M. L. Medus, G. E. Gomez, L. F. Zacchi, P. M. Couto, C. A. Labriola, M. S. Labanda, R. C. Bielsa, E. M. Clérico, B. L. Schulz, J. J. Caramelo, N-glycosylation Triggers a Dual Selection Pressure in Eukaryotic Secretory Proteins. Sci. Rep. 7, 8788 (2017).

35. M. F. Holick, A. Shirvani, N. Charoenngam, Fetal Fractures in an Infant with Maternal Ehlers-Danlos Syndrome, CCDC134 Pathogenic Mutation and a Negative Genetic Test for Osteogenesis Imperfecta. Children **8** (2021).

36. T. M. Ali, B. D. W. Linnenkamp, G. L. Yamamoto, R. S. Honjo, H. Cabral de Menezes Filho, C. A. Kim, D. R. Bertola, The recurrent homozygous translation start site variant in CCDC134 in an individual with severe osteogenesis imperfecta of non-Morrocan ancestry. Am. J. Med. Genet. A 188, 1545–1549 (2022).

37. M. Jovanovic, G. Guterman-Ram, J. C. Marini, Osteogenesis Imperfecta: Mechanisms and Signaling Pathways Connecting Classical and Rare OI Types. Endocr. Rev. 43, 61–90 (2022).

38. S. Moosa, G. L. Yamamoto, L. Garbes, K. Keupp, A. Beleza-Meireles, C. A. Moreno, E. R. Valadares, S. B. de Sousa, S. Maia, J. Saraiva, R. S. Honjo, C. A. Kim, H. Cabral de Menezes, E. Lausch, P. V. Lorini, A. Lamounier Jr, T. C. B. Carniero, C. Giunta, M. Rohrbach, M. Janner, O. Semler, F. Beleggia, Y. Li, G. Yigit, N. Reintjes, J. Altmüller, P. Nürnberg, D. P. Cavalcanti, B. Zabel, M. L. Warman, D. R. Bertola, B. Wollnik, C. Netzer, Autosomal-Recessive Mutations in MESD Cause Osteogenesis Imperfecta. Am. J. Hum. Genet. 105, 836–843 (2019).

39. C. M. Laine, K. S. Joeng, P. M. Campeau, R. Kiviranta, K. Tarkkonen, M. Grover, J. T. Lu, M. Pekkinen, M. Wessman, T. J. Heino, V. Nieminen-Pihala, M. Aronen, T. Laine, H. Kröger, W. G. Cole, A.-E. Lehesjoki, L. Nevarez, D. Krakow, C. J. R. Curry, D. H. Cohn, R. A. Gibbs, B. H. Lee, O. Mäkitie, WNT1 mutations in early-onset osteoporosis and osteogenesis imperfecta. N. Engl. J. Med. 368, 1809–1816 (2013).

40. K. Keupp, F. Beleggia, H. Kayserili, A. M. Barnes, M. Steiner, O. Semler, B. Fischer, G. Yigit, C. Y. Janda, J. Becker, S. Breer, U. Altunoglu, J. Grünhagen, P. Krawitz, J. Hecht, T. Schinke, E. Makareeva, E. Lausch, T. Cankaya, J. A. Caparrós-Martín, P. Lapunzina, S. Temtamy, M. Aglan, B. Zabel, P. Eysel, F. Koerber, S. Leikin, K. C. Garcia, C. Netzer, E. Schönau, V. L. Ruiz-Perez, S. Mundlos, M. Amling, U. Kornak, J. Marini, B. Wollnik, Mutations in WNT1 cause different forms of bone fragility. Am. J. Hum. Genet. 92, 565–574 (2013).

41. S. M. Pyott, T. T. Tran, D. F. Leistritz, M. G. Pepin, N. J. Mendelsohn, R. T. Temme, B. A. Fernandez, S. M. Elsayed, E. Elsobky, I. Verma, S. Nair, E. H. Turner, J. D. Smith, G. P. Jarvik, P. H. Byers, WNT1 mutations in families affected by moderately severe and progressive recessive osteogenesis imperfecta. Am. J. Hum. Genet. 92, 590–597 (2013).

42. S. Fahiminiya, J. Majewski, J. Mort, P. Moffatt, F. H. Glorieux, F. Rauch, Mutations in WNT1 are a cause of osteogenesis imperfecta. J. Med. Genet. 50, 345–348 (2013).

43. L. M. Boyden, J. Mao, J. Belsky, L. Mitzner, A. Farhi, M. A. Mitnick, D. Wu, K. Insogna, R. P. Lifton, High bone density due to a mutation in LDL-receptor-related protein 5. N. Engl. J. Med. 346, 1513– 1521 (2002).

44. Y. Gong, R. B. Slee, N. Fukai, G. Rawadi, S. Roman-Roman, A. M. Reginato, H. Wang, T. Cundy, F. H. Glorieux, D. Lev, M. Zacharin, K. Oexle, J. Marcelino, W. Suwairi, S. Heeger, G. Sabatakos, S. Apte, W. N. Adkins, J. Allgrove, M. Arslan-Kirchner, J. A. Batch, P. Beighton, G. C. Black, R. G. Boles, L. M. Boon, C. Borrone, H. G. Brunner, G. F. Carle, B. Dallapiccola, A. De Paepe, B. Floege, M. L. Halfhide, B. Hall, R. C. Hennekam, T. Hirose, A. Jans, H. Jüppner, C. A. Kim, K. Keppler- Noreuil, A. Kohlschuetter, D. LaCombe, M. Lambert, E. Lemyre, T. Letteboer, L. Peltonen, R. S. Ramesar, M. Romanengo, H. Somer, E. Steichen-Gersdorf, B. Steinmann, B. Sullivan, A. Superti- Furga, W. Swoboda, M. J. van den Boogaard, W. Van Hul, M. Vikkula, M. Votruba, B. Zabel, T. Garcia, R. Baron, B. R. Olsen, M. L. Warman, Osteoporosis-Pseudoglioma Syndrome Collaborative Group, LDL receptor-related protein 5 (LRP5) affects bone accrual and eye development. Cell **107**, 513–523 (2001).

45. M. Kato, M. S. Patel, R. Levasseur, I. Lobov, B. H.-J. Chang, D. A. Glass 2nd, C. Hartmann, L. Li, T.-H. Hwang, C. F. Brayton, R. A. Lang, G. Karsenty, L. Chan, Cbfa1-independent decrease in osteoblast proliferation, osteopenia, and persistent embryonic eye vascularization in mice deficient in Lrp5, a Wnt coreceptor. J. Cell Biol. 157, 303–314 (2002).

46. S. J. Rodda, A. P. McMahon, Distinct roles for Hedgehog and canonical Wnt signaling in specification, differentiation and maintenance of osteoblast progenitors. Development 133, 3231– 3244 (2006).

47. T. P. Hill, D. Später, M. M. Taketo, W. Birchmeier, C. Hartmann, Canonical Wnt/beta-catenin signaling prevents osteoblasts from differentiating into chondrocytes. Dev. Cell 8, 727–738 (2005).

48. T. F. Day, X. Guo, L. Garrett-Beal, Y. Yang, Wnt/beta-catenin signaling in mesenchymal progenitors controls osteoblast and chondrocyte differentiation during vertebrate skeletogenesis. Dev. Cell 8, 739–750 (2005).

49. H. Hu, M. J. Hilton, X. Tu, K. Yu, D. M. Ornitz, F. Long, Sequential roles of Hedgehog and Wnt signaling in osteoblast development. Development 132, 49–60 (2005).

50. DepMap: The Cancer Dependency Map Project at Broad Institute. https://depmap.org/portal/.

51. N. A. Cherepanova, R. Gilmore, Mammalian cells lacking either the cotranslational or posttranslocational oligosaccharyltransferase complex display substrate-dependent defects in asparagine linked glycosylation. Sci. Rep. 6, 20946 (2016).

52. S. Shrimal, R. Gilmore, Oligosaccharyltransferase structures provide novel insight into the mechanism of asparagine-linked glycosylation in prokaryotic and eukaryotic cells. Glycobiology 29, 288–297 (2019).

53. M. Napiórkowska, J. Boilevin, T. Sovdat, T. Darbre, J.-L. Reymond, M. Aebi, K. P. Locher, Molecular basis of lipid-linked oligosaccharide recognition and processing by bacterial oligosaccharyltransferase. Nat. Struct. Mol. Biol. 24, 1100–1106 (2017).

54. S. Gerber, C. Lizak, G. Michaud, M. Bucher, T. Darbre, M. Aebi, J.-L. Reymond, K. P. Locher, Mechanism of bacterial oligosaccharyltransferase: in vitro quantification of sequon binding and catalysis. J. Biol. Chem. 288, 8849–8861 (2013).

55. Q. Yan, W. J. Lennarz, Studies on the function of oligosaccharyl transferase subunits. Stt3p is directly involved in the glycosylation process. J. Biol. Chem. 277, 47692–47700 (2002).

56. C. Lizak, S. Gerber, S. Numao, M. Aebi, K. P. Locher, X-ray structure of a bacterial oligosaccharyltransferase. Nature 474, 350–355 (2011).

57. G. Li, Q. Yan, A. Nita-Lazar, R. S. Haltiwanger, W. J. Lennarz, Studies on the N-glycosylation of the subunits of oligosaccharyl transferase in Saccharomyces cerevisiae. J. Biol. Chem. 280, 1864–1871 (2005).

58. J. D. Huck, N. L. Que, F. Hong, Z. Li, D. T. Gewirth, Structural and Functional Analysis of GRP94 in the Closed State Reveals an Essential Role for the Pre-N Domain and a Potential Client-Binding Site. Cell Rep. 20, 2800–2809 (2017).

59. R. Kornfeld, S. Kornfeld, Assembly of asparagine-linked oligosaccharides. Annu. Rev. Biochem. 54, 631–664 (1985).

60. A. Sharma, M. Mariappan, S. Appathurai, R. S. Hegde, “In Vitro Dissection of Protein Translocation into the Mammalian Endoplasmic Reticulum” in Protein Secretion: Methods and Protocols, A. Economou, Ed. (Humana Press, Totowa, NJ, 2010), pp. 339–363.

61. A. Sundaram, M. Yamsek, F. Zhong, Y. Hooda, R. S. Hegde, R. J. Keenan, Substrate-driven assembly of a translocon for multipass membrane proteins. Nature 611, 167–172 (2022).

62. L. Smalinskaitė, M. K. Kim, A. J. O. Lewis, R. J. Keenan, R. S. Hegde, Mechanism of an intramembrane chaperone for multipass membrane proteins. Nature 611, 161–166 (2022).

63. S. Pfeffer, J. Dudek, M. Gogala, S. Schorr, J. Linxweiler, S. Lang, T. Becker, R. Beckmann, R. Zimmermann, F. Förster, Structure of the mammalian oligosaccharyl-transferase complex in the native ER protein translocon. Nat. Commun. 5, 3072 (2014).

64. R. S. Hegde, R. J. Keenan, The mechanisms of integral membrane protein biogenesis. Nat. Rev. Mol. Cell Biol. 23, 107–124 (2022).

65. M. Gemmer, M. L. Chaillet, J. van Loenhout, R. Cuevas Arenas, D. Vismpas, M. Gröllers-Mulderij, F. A. Koh, P. Albanese, R. A. Scheltema, S. C. Howes, A. Kotecha, J. Fedry, F. Förster, Visualization of translation and protein biogenesis at the ER membrane. Nature, 1–8 (2023).

66. M. Marzec, D. Eletto, Y. Argon, GRP94: An HSP90-like protein specialized for protein folding and quality control in the endoplasmic reticulum. Biochim. Biophys. Acta 1823, 774–787 (2012).

67. S. Wanderling, B. B. Simen, O. Ostrovsky, N. T. Ahmed, S. M. Vogen, T. Gidalevitz, Y. Argon, GRP94 is essential for mesoderm induction and muscle development because it regulates insulin- like growth factor secretion. Mol. Biol. Cell 18, 3764–3775 (2007).

68. C. D. Marceau, A. S. Puschnik, K. Majzoub, Y. S. Ooi, S. M. Brewer, G. Fuchs, K. Swaminathan, M. A. Mata, J. E. Elias, P. Sarnow, J. E. Carette, Genetic dissection of Flaviviridae host factors through genome-scale CRISPR screens. Nature 535, 159–163 (2016).

69. J. Henriksson, X. Chen, T. Gomes, U. Ullah, K. B. Meyer, R. Miragaia, G. Duddy, J. Pramanik, K. Yusa, R. Lahesmaa, S. A. Teichmann, Genome-wide CRISPR Screens in T Helper Cells Reveal Pervasive Crosstalk between Activation and Differentiation. Cell 176, 882–896.e18 (2019).

70. D. R. Amici, J. M. Jackson, M. I. Truica, R. S. Smith, S. A. Abdulkadir, M. L. Mendillo, FIREWORKS: a bottom-up approach to integrative coessentiality network analysis. Life Sci Alliance 4 (2021).

## REFERENCES

71. E. Campeau, V. E. Ruhl, F. Rodier, C. L. Smith, B. L. Rahmberg, J. O. Fuss, J. Campisi, P. Yaswen, P. K. Cooper, P. D. Kaufman, A versatile viral system for expression and depletion of proteins in mammalian cells. PLoS One 4, e6529 (2009).

72. G. V. Pusapati, J. H. Kong, B. B. Patel, A. Krishnan, A. Sagner, M. Kinnebrew, J. Briscoe, L. Aravind, R. Rohatgi, CRISPR Screens Uncover Genes that Regulate Target Cell Sensitivity to the Morphogen Sonic Hedgehog. Dev. Cell 44, 113–129.e8 (2018).

73. J. G. Doench, N. Fusi, M. Sullender, M. Hegde, E. W. Vaimberg, K. F. Donovan, I. Smith, Z. Tothova, C. Wilen, R. Orchard, H. W. Virgin, J. Listgarten, D. E. Root, Optimized sgRNA design to maximize activity and minimize off-target effects of CRISPR-Cas9. Nat. Biotechnol. 34, 184–191 (2016).

74. J. Joung, S. Konermann, J. S. Gootenberg, O. O. Abudayyeh, R. J. Platt, M. D. Brigham, N. E. Sanjana, F. Zhang, Genome-scale CRISPR-Cas9 knockout and transcriptional activation screening. Nat. Protoc. 12, 828–863 (2017).

75. W. Li, H. Xu, T. Xiao, L. Cong, M. I. Love, F. Zhang, R. A. Irizarry, J. S. Liu, M. Brown, X. S. Liu, MAGeCK enables robust identification of essential genes from genome-scale CRISPR/Cas9 knockout screens. Genome Biol. 15, 554 (2014).

76. V. S. W. Li, S. S. Ng, P. J. Boersema, T. Y. Low, W. R. Karthaus, J. P. Gerlach, S. Mohammed, A. J. R. Heck, M. M. Maurice, T. Mahmoudi, H. Clevers, Wnt signaling through inhibition of β-catenin degradation in an intact Axin1 complex. Cell 149, 1245–1256 (2012).

77. N. E. Sanjana, O. Shalem, F. Zhang, Improved vectors and genome-wide libraries for CRISPR screening. Nat. Methods 11, 783–784 (2014).

78. A. Bermudez, S. J. Pitteri, Enrichment of Intact Glycopeptides Using Strong Anion Exchange and Electrostatic Repulsion Hydrophilic Interaction Chromatography. Methods Mol. Biol. 2271, 107–120 (2021).

79. E. Shishkova, A. S. Hebert, M. S. Westphall, J. J. Coon, Ultra-High Pressure (>30,000 psi) Packing of Capillary Columns Enhancing Depth of Shotgun Proteomic Analyses. Anal. Chem. 90, 11503– 11508 (2018).

80. N. M. Riley, S. A. Malaker, M. D. Driessen, C. R. Bertozzi, Optimal Dissociation Methods Differ for N- and O-Glycopeptides. J. Proteome Res. 19, 3286–3301 (2020).

81. J. Cox, M. Mann, MaxQuant enables high peptide identification rates, individualized p.p.b.-range mass accuracies and proteome-wide protein quantification. Nat. Biotechnol. 26, 1367–1372 (2008).

82. D. R. Brademan, I. J. Miller, N. W. Kwiecien, D. J. Pagliarini, M. S. Westphall, J. J. Coon, E. Shishkova, Argonaut: A Web Platform for Collaborative Multi-omic Data Visualization and Exploration. Patterns (N Y) 1 (2020).

83. D. A. Polasky, F. Yu, G. C. Teo, A. I. Nesvizhskii, Fast and comprehensive N- and O-glycoproteomics analysis with MSFragger-Glyco. Nat. Methods 17, 1125–1132 (2020).

84. P. Walter, G. Blobel, Preparation of microsomal membranes for cotranslational protein translocation. Methods Enzymol. 96, 84–93 (1983).

